# Ultrasonic emissions of sunflower and buckwheat carry species identity in their spectrum and drought in their rate

**DOI:** 10.64898/2026.11.17.767468

**Authors:** Bedabrata Saha, Md. Intesaful Haque, Marcin Stróżecki, Anshu Rastogi

**Affiliations:** Department of Bioclimatology, Faculty of Environmental and Mechanical Engineering, Poznań, University of Life Sciences, Piątkowska 94, 60-649 Poznań, Poland

**Keywords:** Acoustic emission, chlorophyll fluorescence, drought, *Fagopyrum esculentum*, *Helianthus annuus*, stomatal conductance, xylem embolism

## Abstract

Plants under water deficit produce ultrasonic acoustic emissions that can be recorded in air. Species have been compared mainly by peak frequency. We report, to our knowledge, the first airborne recordings of sunflower (*Helianthus annuus*) and buckwheat (*Fagopyrum esculentum*), with gas exchange and chlorophyll fluorescence on the same plants. Across 119 recordings of 30 min each, made on two drought and two watered end plants per species, every drought plant emitted more events over the series than the watered plants. Rates rose 107 fold in sunflower and 5.3 fold in buckwheat, coincided with low stomatal conductance, and later fell in single plants. In sunflower, emission was already raised when maximum Photosystem II quantum yield was similar between treatments. At equal loudness, sunflower events carried a median of 24% of their energy at 40-80 kHz, against 2% in buckwheat, in drought and watered plants alike, although both species peaked near 30 kHz. In a proof of concept on four plants per species, this share identified a held-out plant’s species in 96% of draws of five events. Airborne ultrasound thus carries species identity in its spectrum and drought in its event rate, two signals a non-contact sensor could combine.

**Highlight:** Drought raised airborne ultrasonic event rates in sunflower and buckwheat. In a proof-of-concept test on eight plants, the share of event energy above 40 kHz identified the species.

## Introduction

Water deficit is a major cause of yield loss in crops, and early signs of plant water stress are useful for irrigation management (Saha and Rastogi, 2026). Acoustic emissions (AE) from the plant itself are one candidate signal. They are produced in the water-conducting tissue as it dries (SharathKumar et al., 2026). In a greenhouse comparison of 10 sensor types on tomato, AE recorded with an ultrasound microphone near the stem, stomatal conductance and stem diameter all changed within two days of withholding water (Dutta et al., 2025), and contact AE sensors detected osmotic stress in hydroponic lettuce before visible wilting (Hase and Ogihara, 2026).

Ultrasonic AE from drying xylem has been studied for decades with sensors attached to the stem (Milburn and Johnson, 1966; Tyree and Dixon, 1983). Many events arise when cavitation breaks the water column under tension, and their acoustic energy increases with conduit size (Ponomarenko et al., 2014). The settling time and peak frequency of ultrasonic pulses were related to xylem vessel radius in 10 species (Dutta et al., 2022). In the field, emissions coincided with stem shrinkage that tracks xylem tension (Hölttä et al., 2005), and in potted wheat events became frequent once the leaves curled (Jia et al., 2006). Some events come from other sources, such as cell wall shrinkage and cell damage late in dehydration (Vergeynst et al., 2016; Lamacque et al., 2022; Boccato et al., 2024; Nardini et al., 2024). Event counts are now used to build vulnerability curves in woody plants (De Boeck and Steppe, 2025). Stomata usually close at higher water potentials than those that start embolism (Martin-StPaul et al., 2017), so emissions would be expected mainly once stomata have closed.

Khait et al. (2023) showed that tomato and tobacco emit airborne ultrasonic sounds when dried or cut. Drought stressed tomato emitted about 35 sounds per hour, watered plants fewer than one and pots with soil alone none. Mean peak frequency differed between the two species, and a classifier separated species and treatments. Bonisoli et al. (2024; 2025) recorded two to three events per 30 min from bean and tomato with the low-cost microphone model used here. Microphones next to the stems of tree saplings detected induced embolism outdoors (Klaminder et al., 2025).

A recent review in this journal described airborne ultrasound as a source of acoustic phenotypes that can indicate plant water status, with xylem cavitation as the main proposed source (Font Bordera et al., 2026). It concluded that emissions are specific to both species and stress, that their acoustic signature probably reflects vessel diameter and tissue anatomy, and that machine-learning classification could support precision agriculture. It also noted that not every emission need come from cavitation, that current datasets are small and from controlled conditions, and that more species are needed. Airborne recordings exist for few crops, and it is unclear whether species differ in the spectral content of their emissions when loudness is matched. Event counts depend on detection thresholds and electrical noise, and protocols are not standardised (SharathKumar et al., 2026). When several plants share one chamber, it is also unclear which plant produced an event, and some authors have questioned how much information airborne plant sounds carry (Son et al., 2024).

Here we recorded drought and watered plants of sunflower (*Helianthus annuus* L.), a tall oilseed crop with a thick stem, and common buckwheat (*Fagopyrum esculentum* Moench), a short-season pseudocereal described as highly sensitive to drought (Hou et al., 2019), with the same chamber, microphones and detection thresholds. To our knowledge, airborne emissions of neither species have been reported. We addressed three points raised by Font Bordera et al. (2026). First, we tested whether the event rate rises under drought and how it relates to substrate drying, stomatal conductance and PSII function measured on the same plants. Second, we compared the spectral content of events of the two species at equal loudness, in drought and watered plants. Third, we tested, as a proof of concept, whether events identify the species of plants not used to train a classifier. We show that the share of event energy at 40-80 kHz differs about 10 fold between the two species, although their events peak at a similar frequency, and that this share identifies the species. Drought raised the event rate, which coincided with low stomatal conductance and fell again in single plants as they dried further.

## Materials and methods

### Plant material and drought treatment

Sunflower (*Helianthus annuus* L.) and common buckwheat (*Fagopyrum esculentum* Moench) were sown individually in 1 L pots (18.77 g empty) filled with 150 g of commercial peat substrate (pH 5.5-6.5; 47.7 g dry mass). Plants grew in a glasshouse at Poznań University of Life Sciences, Poland, under natural light (midday photosynthetic photon flux density 400-1100 µmol m^−2^ s^−1^), at 28-35 °C by day, 15-20 °C at night and 60-70% relative humidity. They were watered to container capacity every two days and fertilised with a complete nutrient solution. At 37 days after sowing, six uniform plants of each species were selected and their pots brought to similar weights. Three became drought plants, and three remained watered controls that were watered every two days until water dripped from the pot.

Watering of the drought plants stopped at 37 days after sowing (DAS). Sunflower was recorded once the plants began to wilt, at 42, 44 and 47 DAS (5-10 days without water), and buckwheat at 44, 48, 51, 55 and 58 DAS (7-21 days without water). Plants were always recorded before watering. After recording, drought pots were rewatered to an estimated 40-50% of field capacity, allowing for the estimated plant mass.

### Pot weights and substrate water

Pots were weighed after recording on each recording day and at 43 DAS (sunflower), and drought pots also after rewatering. Field capacity was determined on three 100 g substrate samples that were saturated, drained until dripping stopped, weighed and dried at 60 °C for 36 h. The substrate held 5.41 ± 0.11 g water g^−1^ dry mass at field capacity, or 258 g per pot. Substrate water (% of field capacity) was calculated as (pot weight − 66.5 g) ÷ 258 g × 100, where 66.5 g is the pot plus dry substrate. Plant fresh mass was not subtracted, so these values are upper limits. Daily water loss of the drought pots was calculated between successive weighing, including the unweighed additions.

### Leaf gas exchange and chlorophyll fluorescence

Leaf gas exchange and chlorophyll fluorescence were measured with a GFS-3000 system with a fluorescence module (Heinz Walz GmbH, Effeltrich, Germany) on fully expanded leaves of the plants used for the recordings, at 44 and 47 DAS for sunflower and at 48 and 51 DAS for buckwheat, between 11:40 and 15:45 h. The measurement program was the same as in Haque et al. (2026). The cuvette was set to a photosynthetic photon flux density of 1000 µmol m^−2^ s^−1^ from the integrated LED source, a reference CO_2_ concentration of 400 µmol mol^−1^, a water vapour concentration of 17,000 µmol mol^−1^ and a flow rate of 750 µmol s^−1^, and leaf temperature followed ambient conditions. Two leaves were measured on each plant, each once after the readings had reached steady state. We recorded stomatal conductance to water vapour (*g*_s_), transpiration (*E*), intercellular CO_2_ concentration (*c*_i_) and the effective quantum yield of PSII (ΦPSII).

Chlorophyll *a* fluorescence transients (OJIP) were recorded with a FluorPen (Photon Systems Instruments, Drásov, Czech Republic) on two leaves of each recorded plant after 20 min of dark adaptation in leaf clips, as in Haque et al. (2026), at 42, 44 and 47 DAS for sunflower and at 44, 51, 55 and 58 DAS for buckwheat. From each transient we took the maximum quantum yield of PSII (*F*_v_/*F*_m_), the performance index on absorption basis (*PI_abs_*) and the relative variable fluorescence at the J (2 ms) and I (30 ms) steps (*V*_J_, *V*_I_) (Strasser et al., 2000). Transients were plotted as fluorescence relative to *F*_o_ and as the difference in relative variable fluorescence between drought and watered plants. At 42 and 44 DAS the FluorPen recorded a background (Bckg) of 139 instead of 422 for some sunflower leaves. These readings had lower minimal and maximal fluorescence, and gave a higher *PI_abs_*and a lower *V*_J_ than the other leaf of the same plant. Fluorescence variables were therefore compared only between leaves with the same background value, so 42 DAS was not tested and 44 DAS had two drought plants. Two drought buckwheat leaves at 51 DAS had an *F*_v_/*F*_m_ of 0.20 and 0.32. Their *F*_o_ lay within the range of the other leaves measured that day, while their *F*_m_ was 2-3 times lower than in any other leaf, so the low values came from a lack of variable fluorescence. Leaves with an *F*_v_/*F*_m_ of 0.47-0.54 also occurred in drought plants at 55 DAS. These readings were retained, and results are also reported without them.

### Recording set-up

Recordings were made in a wooden chamber lined with pyramidal acoustic foam (62×62×90 cm, inner width × depth × height; Fig. 1A) with two ultrasonic microphones (Ultramic 384K BLE, Dodotronic, Castel Gandolfo, Italy) powered by mains USB adapters and recording at 384 kHz and 16 bit. The three plants of one treatment stood in a row between the microphones, with M1 facing the plant at one end and M2 the plant at the other end (Fig. 1B-D; 5 cm from the nearest stem). The layout was the same for both species and treatments. Each end plant was recorded in four consecutive 30-min recordings per day. On most days the drought plants were recorded in the morning (10:10-13:15 h) and the controls in the early afternoon (12:40-16:20 h); Supplementary Table S1 gives the time window of each group. The thresholds were set on empty-chamber recordings.

**Fig. 1.**
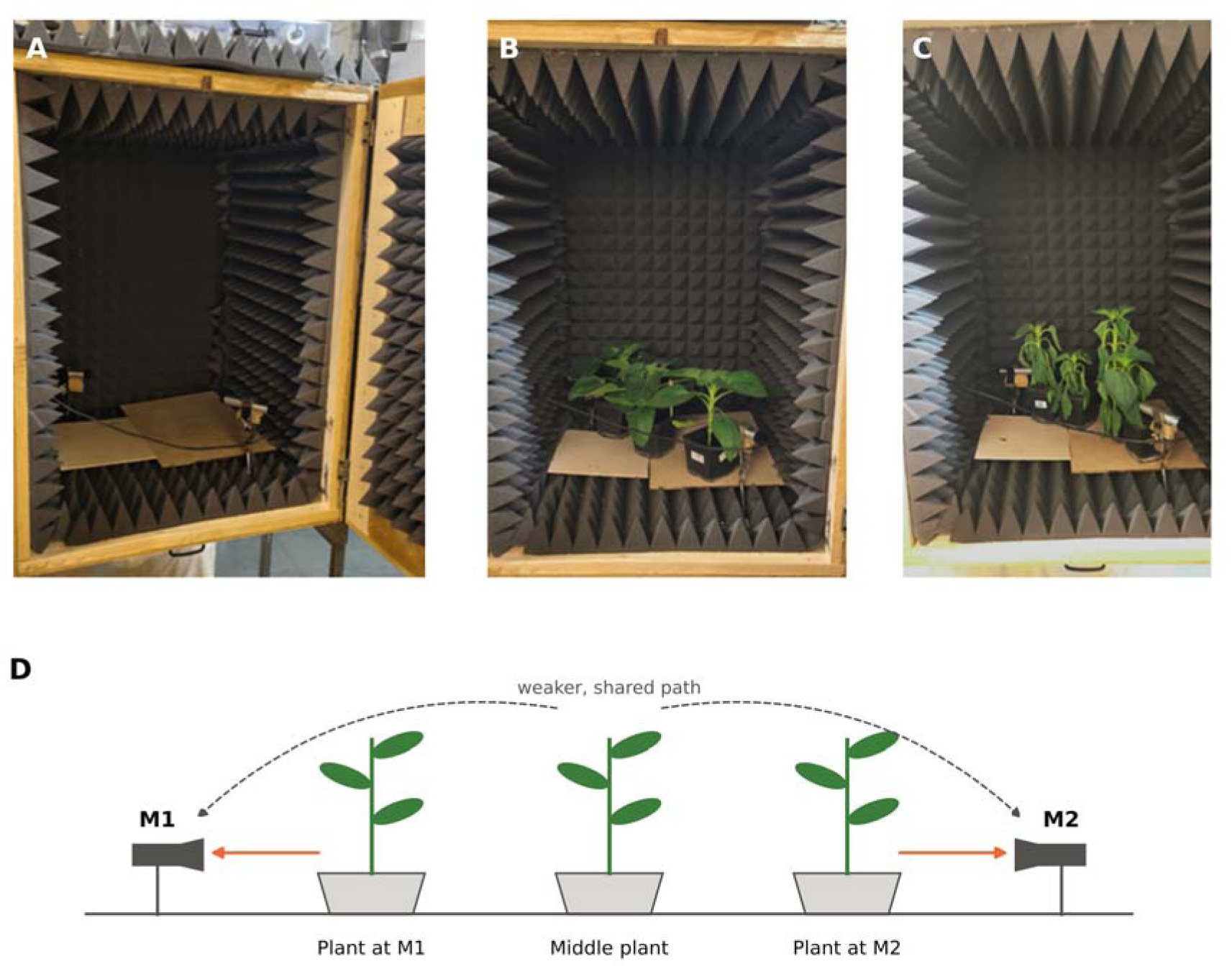
Recording chamber and plant layout. (A) Empty chamber lined with acoustic foam, with the two ultrasonic microphones. (B) Watered and (C) drought plants in a row between the microphones. (D) Layout used in all recordings; orange arrows, direct path from each end plant to its microphone; dashed arrows, path from the middle plant. Not to scale. **Alt text:** Photographs of a foam-lined wooden recording chamber with two microphones, empty, with three upright watered plants and with three wilted drought plants in a row, and a schematic of the plant row between microphones M1 and M2.

### Spike repair and event detection

The mains adapters introduced one-sample electrical spikes, broadband up to 192 kHz (Supplementary Fig. S1). A sample was replaced by the five-point running median when it differed from that median by more than six robust standard deviations and the energy at 150-184 kHz exceeded 9% of that at 20-100 kHz. In six test recordings, 705 of 709 events were detected again after repair, with unchanged peak amplitude (ratio 0.999-1.002).

The repaired signal was split into four bands (20-40, 40-60, 60-80 and 80-100 kHz; fourth-order zero-phase Butterworth filters). Each band threshold was set 1 dB above the highest amplitude in 16 empty-chamber recordings (479 min), giving −50.8, −59.8, −60.7 and −60.5 dBFS (decibels relative to full scale). An event began at a threshold crossing and ended after 10 ms without one. Overlapping events in different bands were merged, and the event level was the highest band peak. A band event was kept when its peak exceeded eight times the robust standard deviation of the band signal within ±50 ms. Recordings with a background above 1.1 times the empty-chamber level were classed as having raised electrical background. Only events above the highest threshold (−50.8 dBFS), called primary events, were analysed, so that every band had the same sensitivity. Events recorded by both microphones were counted once, at the louder microphone, and buckwheat events classed as electrical impulses were removed.

### Waveform check and spectral features

Clips of ±3 ms around each event (3,483 sunflower and 358 buckwheat clips) were spike-repaired and band-pass filtered at 15-150 kHz. The Hilbert envelope gave the event peak and the duration above 25% of the peak. The spectrum was computed over 0.5 ms (Hann window) from 0.1 ms before the peak, minus the spectrum of an equal background window 1.8 ms earlier. From it we took the peak frequency, spectral centroid, bandwidth at −10 dB and the shares of energy at 15-40, 40-80 and 80-150 kHz. The decay time was the time for the 20-40 kHz envelope to fall to 1/e of its peak. Spectrograms used 0.125 ms Hann windows. Buckwheat clips were classed as ringing bursts (≥0.06 ms, stable frequency, most energy below 80 kHz), electrical impulses (<0.04 ms, much energy above 80 kHz) or unclear events. All sunflower recordings had normal background. Species were compared at equal loudness, with primary events between −50.8 and −44 dBFS (915 sunflower events from 37 recordings and 90 buckwheat events from 38 recordings). For these events we also noted the bands in which each event exceeded the threshold, in recordings with normal background.

### Events recorded by both microphones

The microphones ran on separate clocks. For each pair of simultaneous recordings, the clock offset was the most frequent time difference between M1 and M2 events within ±5 s, and a line fitted to the candidate pairs corrected for drift. Event pairs within ±3 ms of this line were classed as shared events. For the shared sunflower events of 42 DAS, the arrival-time difference between events louder at M1 and at M2 was estimated by least squares, and the spectra of the near (louder) and far copies were compared.

### Event size, amplitude index and timing

For the sunflower drought plant at M1, which emitted the most events, the level distribution on each day was described by the share of events above −35 dBFS and by the maximum-likelihood power-law exponent β = n ÷ Σ ln(A_i_ ÷ A_0_), where A_i_ is the peak amplitude of event i and A_0_ that at −50.8 dBFS (a smaller β means more loud events). The within-recording relation between level and the share of energy at 40-80 kHz was estimated by splitting each event level into its recording mean and the deviation from it. As a relative index of emission intensity, the squared peak amplitudes of the primary events, relative to full scale, were summed per 30 min. This summed squared peak amplitude is not calibrated to sound pressure and is not the energy of the whole waveform.

Event timing was tested in recordings with at least 30 primary events, by the coefficient of variation of the intervals between events (1 for random, Poisson, timing) and by the Fano factor of counts in 1, 10 and 60 s bins, compared with 2,000 random placements of the events. Excess short intervals (<0.1, 0.25 and 0.5 s) were tested with a Poisson test. Buckwheat intervals from recordings with at least three events were scaled by their recording mean, pooled and compared with the exponential distribution, and buckwheat counts were tested for overdispersion among the recordings of each plant and day.

### Classification of species and treatment

Logistic regression classifiers were trained on primary events between −50.8 and −44 dBFS, using either the log share of energy at 40-80 kHz alone or eight features (the three energy shares, spectral centroid, peak frequency, bandwidth, duration and decay time), standardised and with balanced class weights. Each of the eight end plants was left out in turn and classified by a model trained on the other seven, and the area under the receiver operating characteristic curve (AUC) was computed from these predictions. To estimate how many events identify a plant, 1-10 events were drawn without replacement 2,000 times from each left-out plant and their predicted log-odds averaged. The smallest plant had 10 events, so the 10 event draws of the two smallest plants contained the same or nearly the same events. Drought and watered plants were classified in the same way within each species and compared with the AUC of the event rate of single plant-days (mean of the four recordings).

### Statistical analysis

The end plant was the experimental unit for treatment effects on emission, with two drought and two watered end plants per species. The four consecutive recordings of a plant on one day were repeated measurements of that plant-day and were not treated as independent replicates. With two plants per treatment, the model-based confidence intervals and p values are approximate, and the treatment comparisons are exploratory. Event counts of each species were analysed with a generalised linear mixed model (log link; glmmTMB) with treatment, recording day and treatment × day as fixed effects, random intercepts for plant and plant-day, and an offset for recording length. A negative binomial model was used for sunflower and a Poisson model for buckwheat, whose counts were not overdispersed, without the 58 DAS recordings, which had no controls. Day-specific rate ratios came from linear combinations of the coefficients with Wald 95% confidence intervals (CI), and the treatment × day interaction was tested with a likelihood-ratio test. The buckwheat model was refitted without the 55 DAS recordings, with normal-background recordings only and with the background ratio as a covariate. At plant level, the mean rate of each end plant over the days with controls was compared between treatments with an exact permutation test stratified by species (36 allocations). Recording background and repaired spike samples were compared between treatments with Mann-Whitney U tests on recordings.

Event levels and spectral features were compared with linear mixed models with plant and recording as random effects and event level as a covariate, with log-transformed energy share at 40-80 kHz and decay time and with Kenward-Roger degrees of freedom. Treatment and species × treatment terms were tested in the same models. Band detection was compared between species with plants as units, as percentage-point differences with 95% CI from 5,000 bootstrap samples of plants within species and permutation tests over the 70 allocations of the eight end plants. For plant-level comparisons, events recorded at each microphone were attributed to the nearest end plant. Gas-exchange and fluorescence variables were averaged over the two leaves of each plant and compared between treatments on each day with two-sided Welch’s t-tests (n = 3 plants). Plants were compared with exact Mann-Whitney U tests, and near and far copies of shared events with Wilcoxon signed-rank tests. All tests were two-sided. Detection was run in R[4.6.2] (tuneR, signal, seewave, readr, dplyr, stringr), mixed models in R (glmmTMB, lme4, lmerTest) and all other analyses in Python 3 (NumPy, SciPy, statsmodels, scikit-learn).

## Results

### Recording quality and event checks

All sunflower recordings had a background of 1.00-1.09 times the empty-chamber level, and drought and watered recordings did not differ in the number of repaired spike samples (p = 0.13 in sunflower and 0.12 in buckwheat). In buckwheat, all recordings at 55 and 58 DAS and the M2 recordings at 44 DAS had a raised background (1.10-1.33 times). Buckwheat events above −50.8 dBFS were mostly ringing bursts with a spectral peak near 28-30 kHz, and weaker events were mostly background peaks or single impulses. The final data set held 1,698 primary events from drought and 198 from watered sunflower, and 72 from drought and 26 from watered buckwheat.

### Drought raised event rates in both species

Each end plant was followed over the recording days (Fig. 2A-D; Table 1). At 42 DAS, both sunflower drought end plants emitted far more events than the controls (103 and 52 against 2.0 and 0.25 events per 30 min). The two drought end plants then followed different courses.

**Fig. 2.**
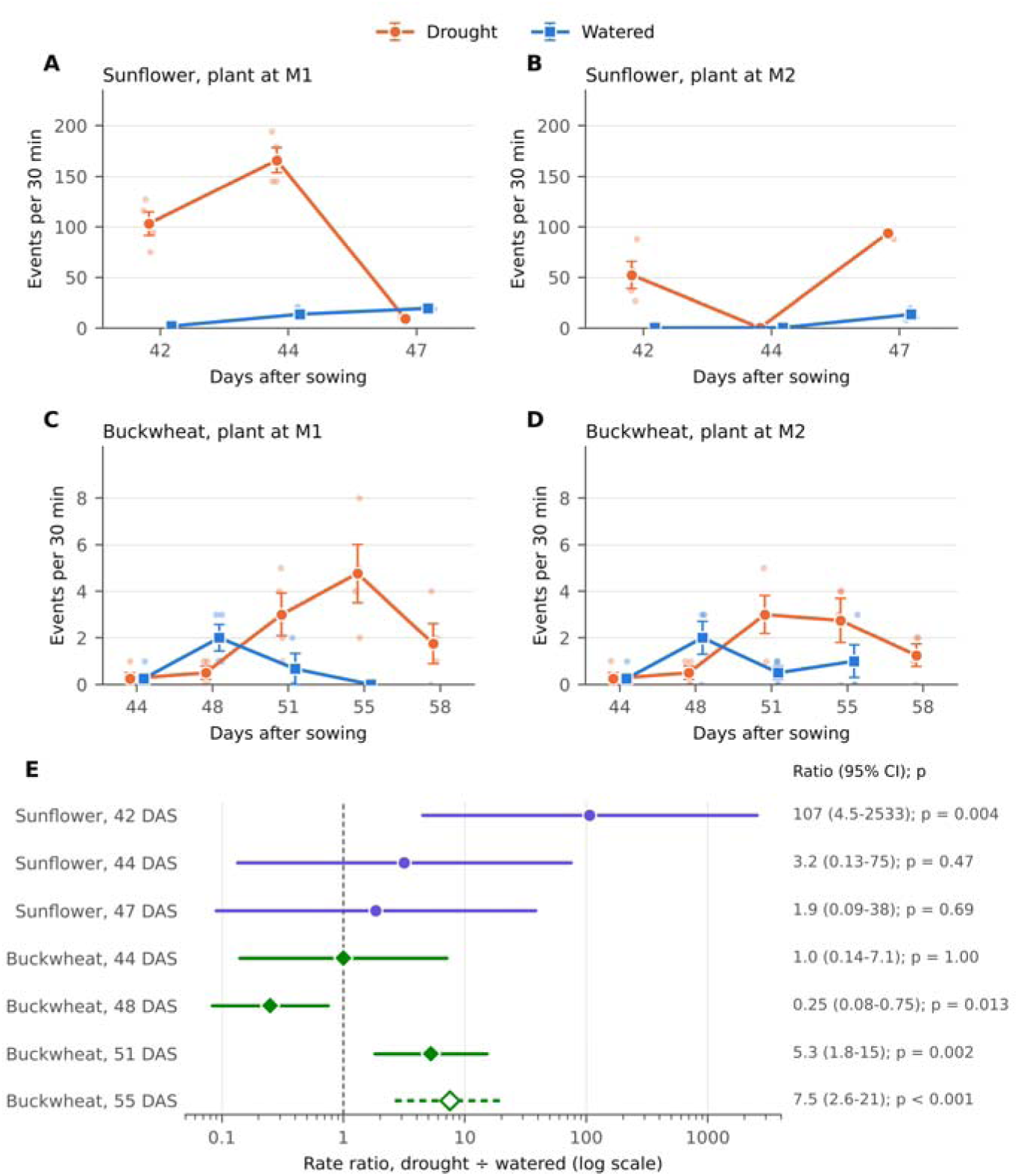
Drought increased ultrasonic event rates in sunflower and buckwheat. (A-D) Each panel follows one drought and one watered end plant over the recording days. Large symbols, mean ± SE of four 30-min recordings (three for the watered buckwheat plant at M1 at 51 DAS); small dots, single recordings; watered buckwheat was not recorded at 58 DAS. (E) Rate ratios with 95% CI from generalised linear mixed models with plant and plant-day as random effects (sunflower, negative binomial, 12 plant-days; buckwheat, Poisson, 16 plant-days); open symbol, raised electrical background in all recordings. M1 and M2, microphones; DAS, days after sowing. **Alt text:** Four dot plots of event rates per 30 minutes over the recording days, showing higher rates in drought than watered plants of both species, and a plot of rate ratios with confidence intervals for each day.

**Table 1.**
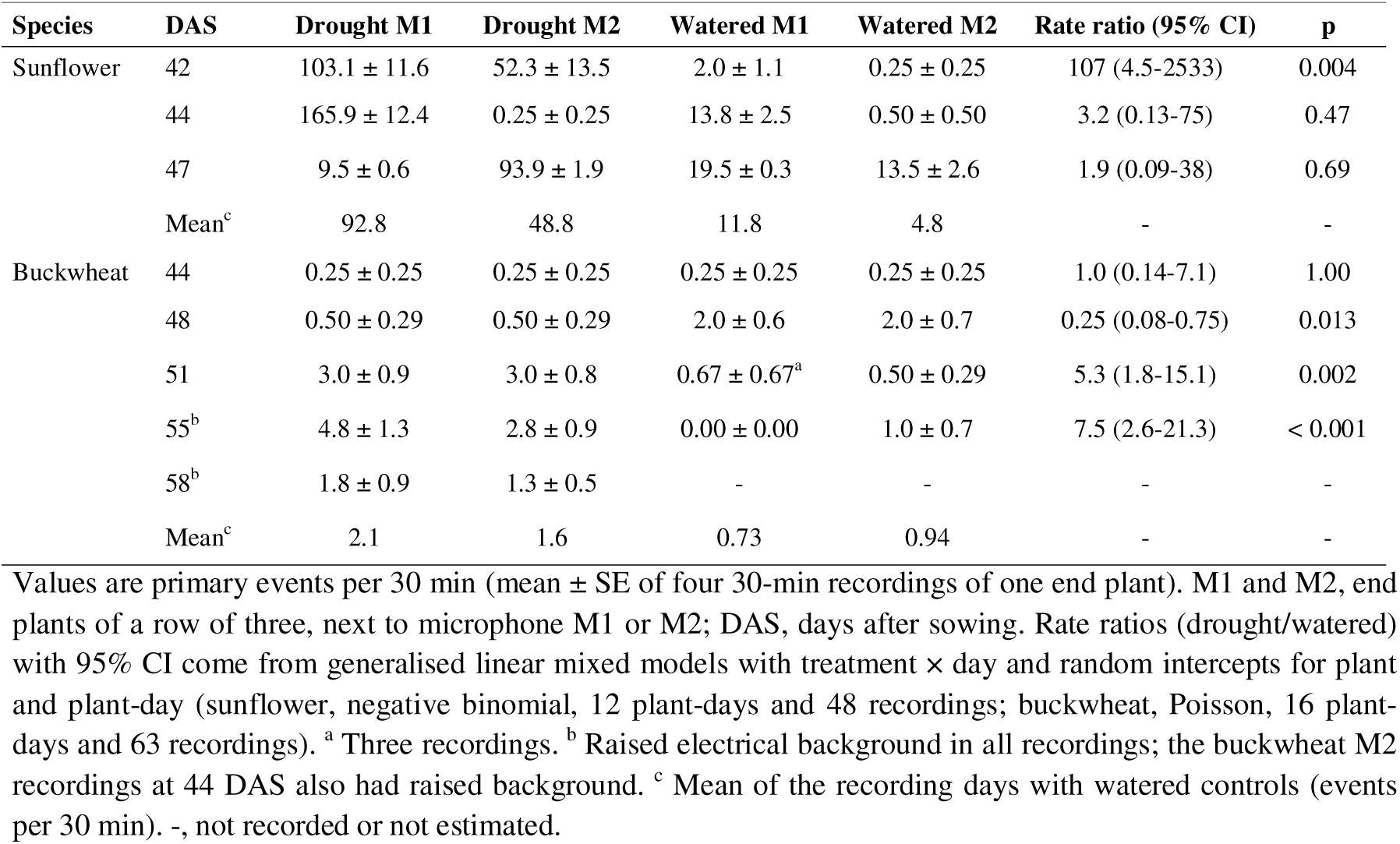
Ultrasonic event rates of sunflower and buckwheat drought plants and watered controls by recording day and microphone position.

The plant at M1 emitted 166 events per 30 min at 44 DAS and 9.5 at 47 DAS (Fig. 2A), and the plant at M2 emitted 0.25 and 94 (Fig. 2B). The control at M2 was also nearly silent at 44 DAS (0.5 events per 30 min), and control rates rose to 13.5-19.5 events per 30 min at 47 DAS.

Buckwheat emitted far fewer events (Fig. 2C, D). Drought and watered plants both emitted 0.25 events per 30 min at 44 DAS, and controls emitted more at 48 DAS (2.0 against 0.5 events per 30 min, 8 against 2 events). Both drought end plants then rose to 3.0 events per 30 min at 51 DAS (controls 0.5-0.67) and to 2.8-4.8 at 55 DAS (controls 0-1.0), and fell to 1.3-1.8 at 58 DAS, when no controls were recorded.

In the mixed model, sunflower drought plants emitted 107 times more events than controls at 42 DAS (95% CI 4.5-2533, p = 0.004; Fig. 2E; Table 1). At 44 and 47 DAS, when only one drought end plant exceeded the controls, the effect was not significant (rate ratios 3.2 and 1.9, p = 0.47 and 0.69). In buckwheat, controls emitted more at 48 DAS (rate ratio 0.25, 95% CI 0.08-0.75, p = 0.013), and drought plants emitted more at 51 DAS (5.3, 1.8-15.1, p = 0.002) and 55 DAS (7.5, 2.6-21.3, p < 0.001). Over the days with controls, both drought end plants of each species emitted more events than both watered end plants (sunflower 93 and 49 against 12 and 4.8 events per 30 min; buckwheat 2.1 and 1.6 against 0.7 and 0.9; Table 1). With two plants per treatment, this complete separation in both species is the most extreme ordering possible (exact permutation test, p = 0.056).

The buckwheat result did not depend on the recordings with raised background (Supplementary Table S2). Without 55 DAS, drought plants still emitted 5.3 times more events than controls at 51 DAS (95% CI 1.8-15.1, p = 0.002), and normal-background recordings alone gave the same value. The local background check removed most detections at 55 DAS in both treatments, so the result of that day is less certain than that of 51 DAS.

### Sunflower events carried more energy above 40 kHz

Events of drought sunflower had a higher median level than those of watered sunflower (−43.1 against −46.9 dBFS), but with two plants per treatment the difference was not significant (3.0 dB, p = 0.35). Buckwheat events lay close to the counting level in both treatments (median −49.5 and −47.9 dBFS, p = 0.34). Single events of both species were short ringing bursts with most of their energy near 30 kHz (Fig. 3A). At equal loudness, the median spectrum of sunflower stayed high above 40 kHz, and that of buckwheat fell steeply above 35 kHz (Fig. 3B). Sunflower events carried a median of 24% of their energy at 40-80 kHz, against 2% in buckwheat (Fig. 3C; Table 2). With plants and recordings as random effects, this share was 9.5 times higher in sunflower (95% CI 6.1-14.7, p < 0.001). Sunflower events also had a higher spectral centroid, were shorter and decayed faster (Table 2). The median peak frequency was similar (30.2 against 28.8 kHz). Although 30% of sunflower events peaked above 35 kHz, against 4% in buckwheat, the mean peak frequency did not differ significantly (4.7 kHz higher in sunflower, 95% CI −1.4 to 10.8, p = 0.11).

**Fig. 3.**
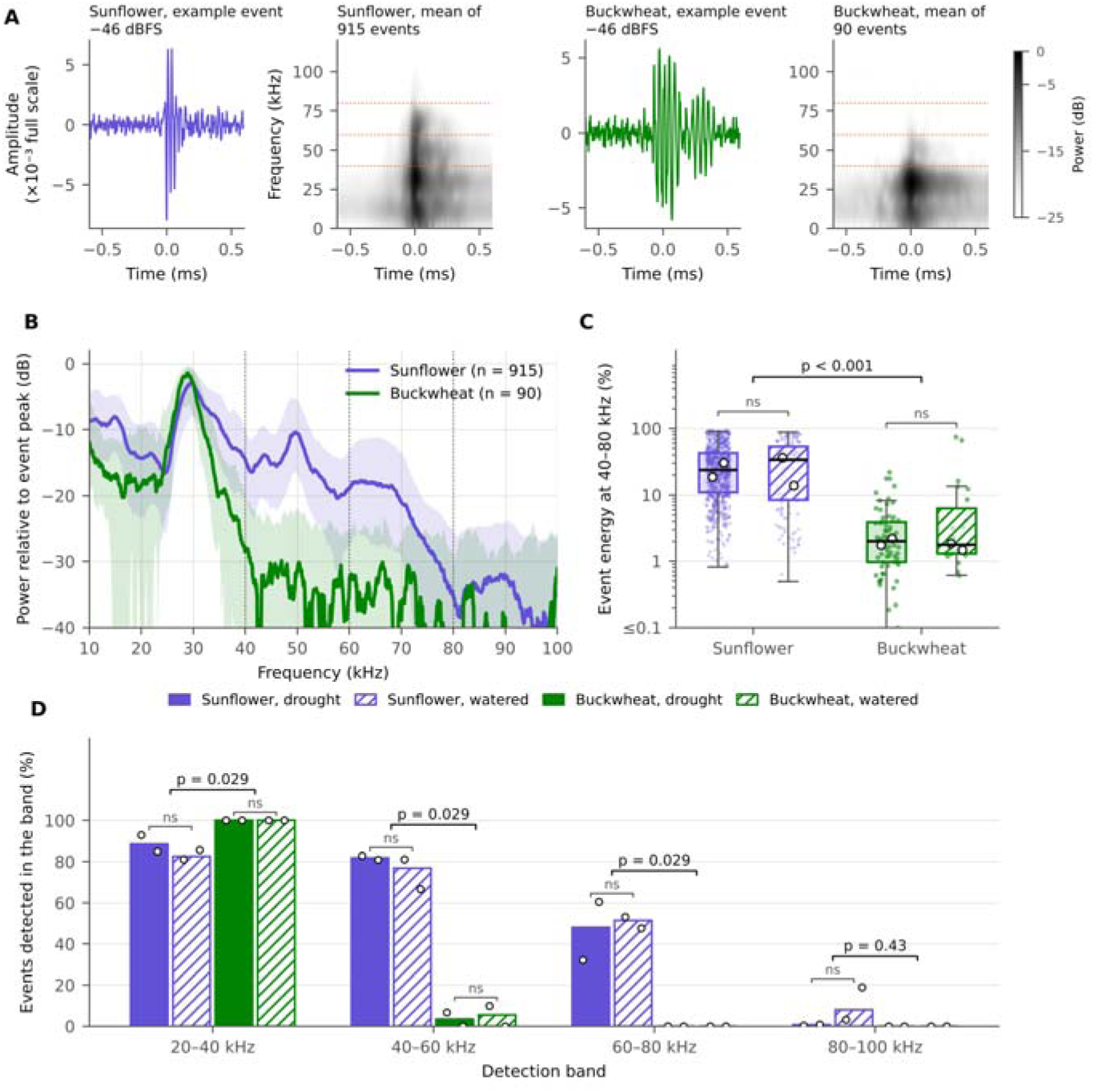
Sunflower events carried more energy above 40 kHz than buckwheat events at equal loudness. Primary events between −50.8 and −44 dBFS from 37 sunflower and 38 buckwheat recordings. (A) Example event and mean normalised spectrogram of each species. (B) Median power spectrum with interquartile range (shading). (C) Boxes, median and interquartile range; whiskers, 1.5 × interquartile range; dots, single events; open circles, plant medians; p, linear mixed models with plant and recording as random effects and event level as covariate. (D) Bars, pooled share of events in recordings with normal background (47 buckwheat events); open circles, single plants; p, two-sided permutation tests with plants as units. Dashed lines, 40, 60 and 80 kHz; ns, p ≥ 0.05. **Alt text:** Example waveforms and mean spectrograms of sunflower and buckwheat events, their median spectra, box plots of the share of energy at 40 to 80 kHz, and bar charts of detection in four frequency bands, all showing more high-frequency energy in sunflower.

**Table 2.** Spectral features and band-specific detection of sunflower and buckwheat events at equal loudness.

| **Feature** | **Sunflower** | **Buckwheat** | **Difference, sunflower − buckwheat (95% CI)** | **p** |
|---|---|---|---|---|
| **Spectral features from raw-waveform clips, all recording days (sunflower n = 915 events from 37 recordings, buckwheat n = 90 events from 38 recordings)** |  |  |  |  |
| Event level (dBFS) | −48.1 (−49.4 to −46.2) | −49.5 (−50.4 to −48.0) | +1.0 dB (0.1 to 1.9) | 0.035 |
| Peak frequency (kHz) | 30.2 (28.6-37.3) | 28.8 (27.2-30.0) | +4.7 kHz (−1.4 to 10.8) | 0.11 |
| Spectral centroid (kHz) | 36.3 (32.8-42.2) | 31.2 (29.3-33.5) | +7.6 kHz (2.6 to 12.5) | 0.009 |
| Bandwidth at −10 dB (kHz) | 7.9 (5.8-10.7) | 6.9 (6.2-8.3) | +0.8 kHz (−0.2 to 1.8) | 0.10 |
| Energy at 15-40 kHz (%) | 73.3 (54.4-86.7) | 95.8 (90.8-97.2) | −27.5 points (−39.0 to −16.0) | < 0.001 |
| Energy at 40-80 kHz (%) | 24.4 (10.8-43.8) | 1.9 (1.1-4.1) | × 9.5 (6.1 to 14.7) | < 0.001 |
| Energy at 80-150 kHz (%) | 1.5 (0.8-2.6) | 2.2 (1.2-4.2) | +0.5 points (−1.7 to 2.8) | 0.60 |
| Duration above 25% of peak (ms) | 0.06 (0.04-0.08) | 0.08 (0.06-0.12) | −0.04 ms (−0.06 to −0.03) | < 0.001 |
| Decay time of 20-40 kHz ringing to 1/e (ms) | 0.07 (0.05-0.10) | 0.12 (0.08-0.15) | × 0.65 (0.57 to 0.74) | < 0.001 |
| **Band-specific detection in all recordings with normal background (sunflower n = 915 events from 37 recordings, buckwheat n = 47 events from 23 recordings)** |  |  |  |  |
| Event level (dBFS) | −48.1 (−49.4 to −46.2) | −49.6 (−50.5 to −48.4) | +1.3 dB (0.4 to 2.1) | 0.006 |
| Events detected in the 20-40 kHz band (%) | 87.7 | 100 | −12.3 points (−17.4 to −7.8) | 0.029 |
| Events detected in the 40-60 kHz band (%) | 81.1 | 4.3 | +76.8 points (70.6 to 81.4) | 0.029 |
| Events detected in the 60-80 kHz band (%) | 48.5 | 0 | +48.5 points (33.9 to 60.1) | 0.029 |
| Events detected in the 80-100 kHz band (%) | 1.7 | 0 | +1.7 points (0.5 to 8.0) | 0.43 |
| Events strongest in a band above 40 kHz (%) | 18.7 | 0 | +18.7 points (12.9 to 29.0) | 0.029 |

Values are medians (interquartile range) or percentages of primary events between −50.8 and −44 dBFS. Differences in event level and spectral features come from linear mixed models with plant and recording as random effects and event level as a covariate, with Kenward-Roger degrees of freedom; × marks ratios for log-transformed features. Differences in band detection are percentage points, with 95% CI from bootstrap samples of plants within species and two-sided permutation p values with plants as units (70 allocations).

The difference in the share at 40-80 kHz held for every plant and recording day (Fig. 3C). Plant medians ranged from 13.9 to 36.3% in sunflower and from 1.5 to 2.2% in buckwheat (exact Mann-Whitney test, four plants per species, p = 0.029), and all nine sunflower plant-days with at least five events exceeded all seven buckwheat plant-days. Peak frequency did not separate the plants (plant medians 29.1-31.7 against 28.2-29.5 kHz, p = 0.11). Within each species the share did not differ between drought and watered plants (sunflower 23.5% against 33.6%, p = 0.67; buckwheat 2.0% against 1.8%, p = 0.33; species × treatment, p = 0.20). The difference also held in the narrowest level class (30% against 2% between −50.8 and −49.5 dBFS). Across all single events, the share at 40-80 kHz separated the species with an AUC of 0.92, against 0.67 for peak frequency.

The detection bands showed the same difference (Fig. 3D; Table 2). Of 915 sunflower events between −50.8 and −44 dBFS, 81% were detected in the 40-60 kHz band and 49% in the 60-80 kHz band. Of 47 buckwheat events from recordings with normal background, 4% reached the 40-60 kHz band and none reached higher bands. Detection at 40-60 kHz was 77 percentage points higher in sunflower (95% CI 71-81 from resampling plants), and every sunflower plant exceeded every buckwheat plant in each band measure (plant-level permutation test, p = 0.029, the lowest possible with four plants per species; Supplementary Table S3). In sunflower, detection at 40-60 kHz was similar in drought and watered plants (81-83% against 67-81%), and the 80-100 kHz band was reached by 3% and 19% of events of the two watered plants, against 0.3% and 0.9% of the drought plants.

### Events recorded by both microphones

Shared events were rare. At 42 DAS, 35 events of the sunflower drought plants were recorded by both microphones, 6% of the 621 primary events of that day. Each was 12-27 dB louder at one microphone (Fig. 4A), more than the other shared events (median 20.7 against 5.7 dB, p < 0.001). After correction for clock offset and drift, the two groups differed in arrival time by 2.24 ms (95% CI 2.20-2.27), and single events lay within 0.1 ms of their group median (Fig. 4B). Only six other shared events were found in sunflower and three in buckwheat. The far copy of an event kept its share of energy at 40-80 kHz when the event was louder at M2 (median 81% at both microphones, n = 21), and lost more than half of it when the event was louder at M1 (9.1% against 2.3%, p = 0.011, n = 14; Fig. 4C).

**Fig. 4.**
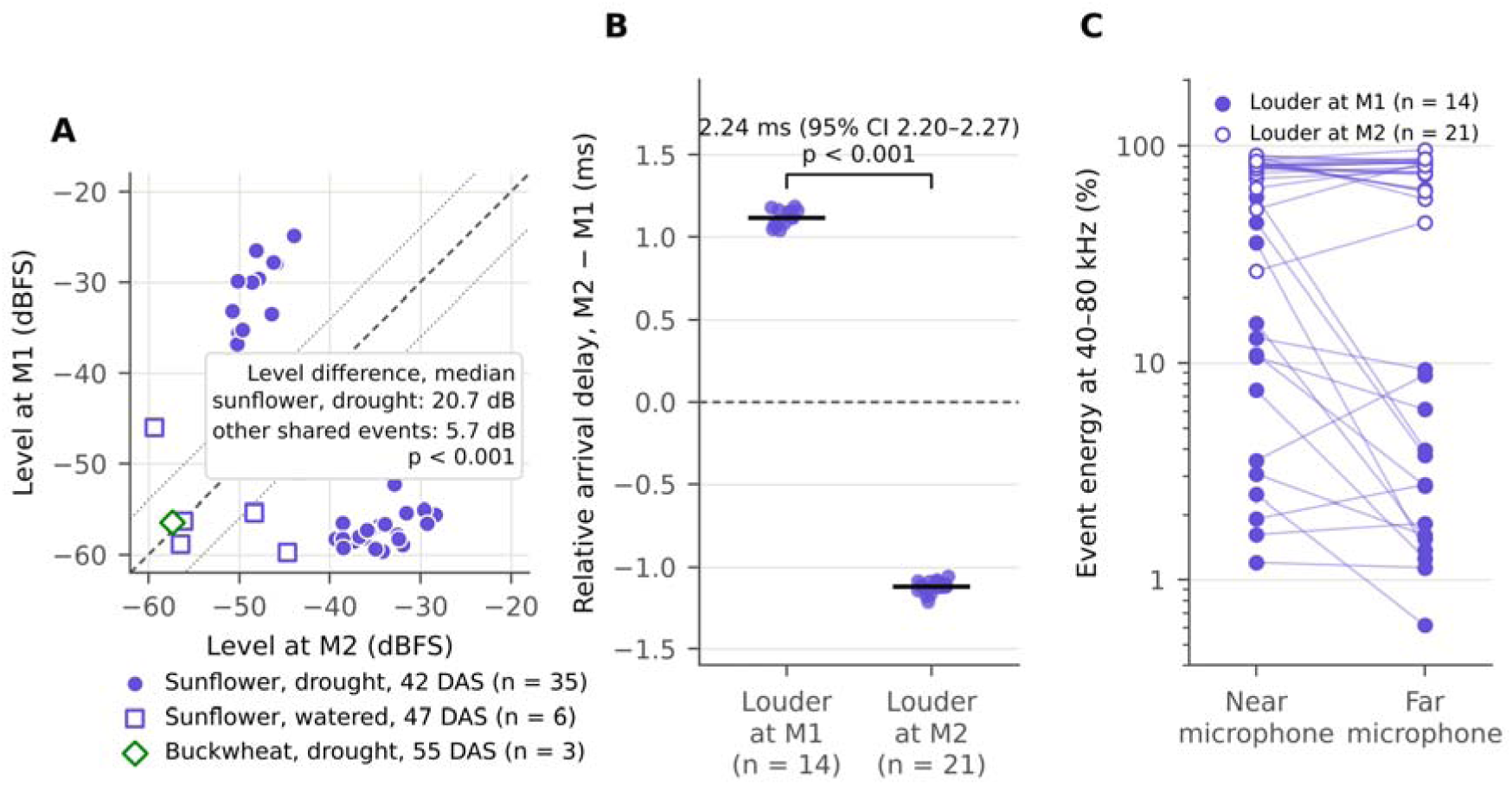
Events recorded by both microphones were much louder at one of them. (A) Level of each shared event at microphones M1 and M2; dashed line, equal levels; dotted lines, 6 dB difference; p, two-sided Mann-Whitney U test. (B) Arrival delay at M2 relative to M1 of the shared sunflower drought events, after removal of clock offset and drift; bars, medians; bracket, difference with 95% CI. (C) Share of energy at 40-80 kHz of the same events at the near (louder) and far microphone; lines join the two copies of each event. DAS, days after sowing. **Alt text:** A scatter plot of event levels at the two microphones with points far from the line of equal level, a dot plot of arrival delays in two separate groups, and paired lines comparing high-frequency energy at the near and far microphone.

### Substrate drying in the drought pots

The drought pots lost water steadily (Fig. 5A, B; Supplementary Table S4). Before rewatering, the sunflower drought pots held at most 50-61% of field capacity at 42 DAS and 36-38% at 47 DAS. At 44 DAS the pots of the end plants at M1 and M2 held at most 49% and 46% of field capacity, and by 47 DAS both had lost 63-64 g of water over three days. The buckwheat drought pots held at most 72-78% of field capacity at 44 DAS, 51-55% at 51 DAS and 34-38% at 58 DAS. Their event rates rose to 3.0-4.8 per 30 min at 42-55% of field capacity (51 and 55 DAS) and fell to 1.3-1.8 at 34-38% (58 DAS).

**Fig. 5.**
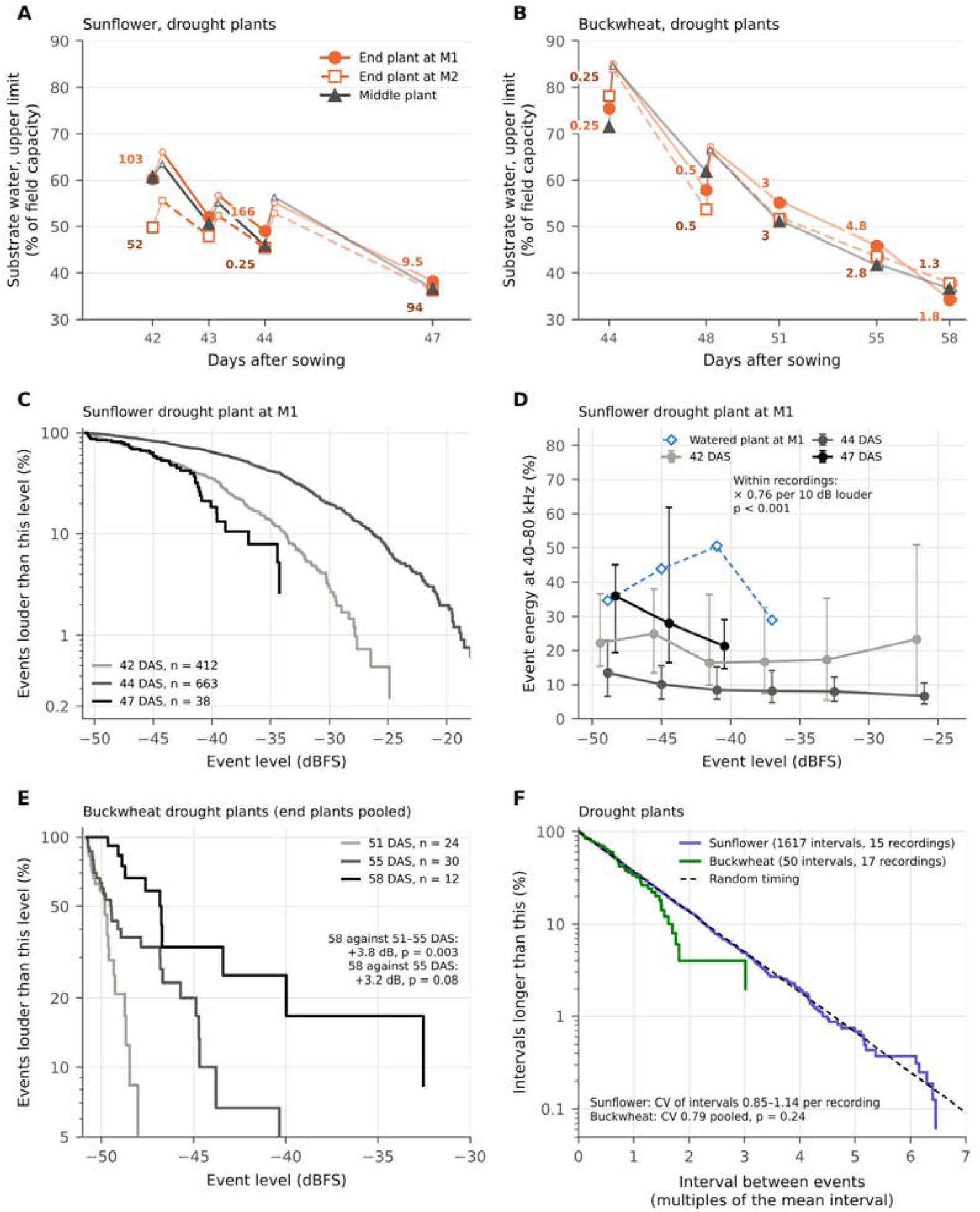
Substrate drying, event size, spectrum and timing in drought plants. (A, B) Substrate water of the three drought pots; small open symbols, after rewatering; numbers, mean events per 30 min of the end plants. (C) Share of events louder than a given level. (D) Median share of energy at 40-80 kHz (interquartile range) in level classes with at least six events; the within-recording factor comes from a linear mixed model. (E) Level distribution of buckwheat drought events; recordings at 55 and 58 DAS had raised electrical background; level differences from linear mixed models. (F) Intervals between events relative to the mean interval of each recording, for recordings with at least 30 (sunflower) or three (buckwheat) events; dashed line, exponential distribution expected for random timing. M1 and M2, microphones; CV, coefficient of variation; p, two-sided Kolmogorov-Smirnov test; DAS, days after sowing. **Alt text:** Line plots of substrate water falling over the recording days in both species, curves of event level distributions and high-frequency energy by level for one sunflower plant, level distributions of buckwheat events on three days, and interval distributions close to the random expectation.

### Stomatal conductance and PSII function of drought plants

Drought plants of both species had much lower stomatal conductance and transpiration than watered plants (Fig. 6A, B; Supplementary Table S5). In sunflower, *g*_s_ of drought plants was 33 and 17 mmol m^−2^ s^−1^ at 44 and 47 DAS, against 237 and 235 mmol m^−2^ s^−1^ in watered plants (p < 0.001 and p = 0.014), and their *c*_i_ rose to 477 µmol mol^−1^ at 47 DAS. In buckwheat, *g*_s_ of drought plants fell from 55 mmol m^−2^ s^−1^ at 48 DAS to 21 mmol m^−2^ s^−1^ at 51 DAS, against 179 and 132 mmol m^−2^ s^−1^ in watered plants (p < 0.001 and p = 0.019). Within each day, the two sunflower drought end plants had similar *g*_s_ (30 and 25 mmol m^−2^ s^−1^ at 44 DAS, 19 and 17 mmol m^−2^ s^−1^ at 47 DAS), although their event rates differed widely (166 against 0.25 and 9.5 against 94 events per 30 min; Supplementary Table S6). ΦPSII differed between treatments only in buckwheat at 48 DAS (0.17 in drought against 0.07 in watered plants, p = 0.039).

**Fig. 6.**
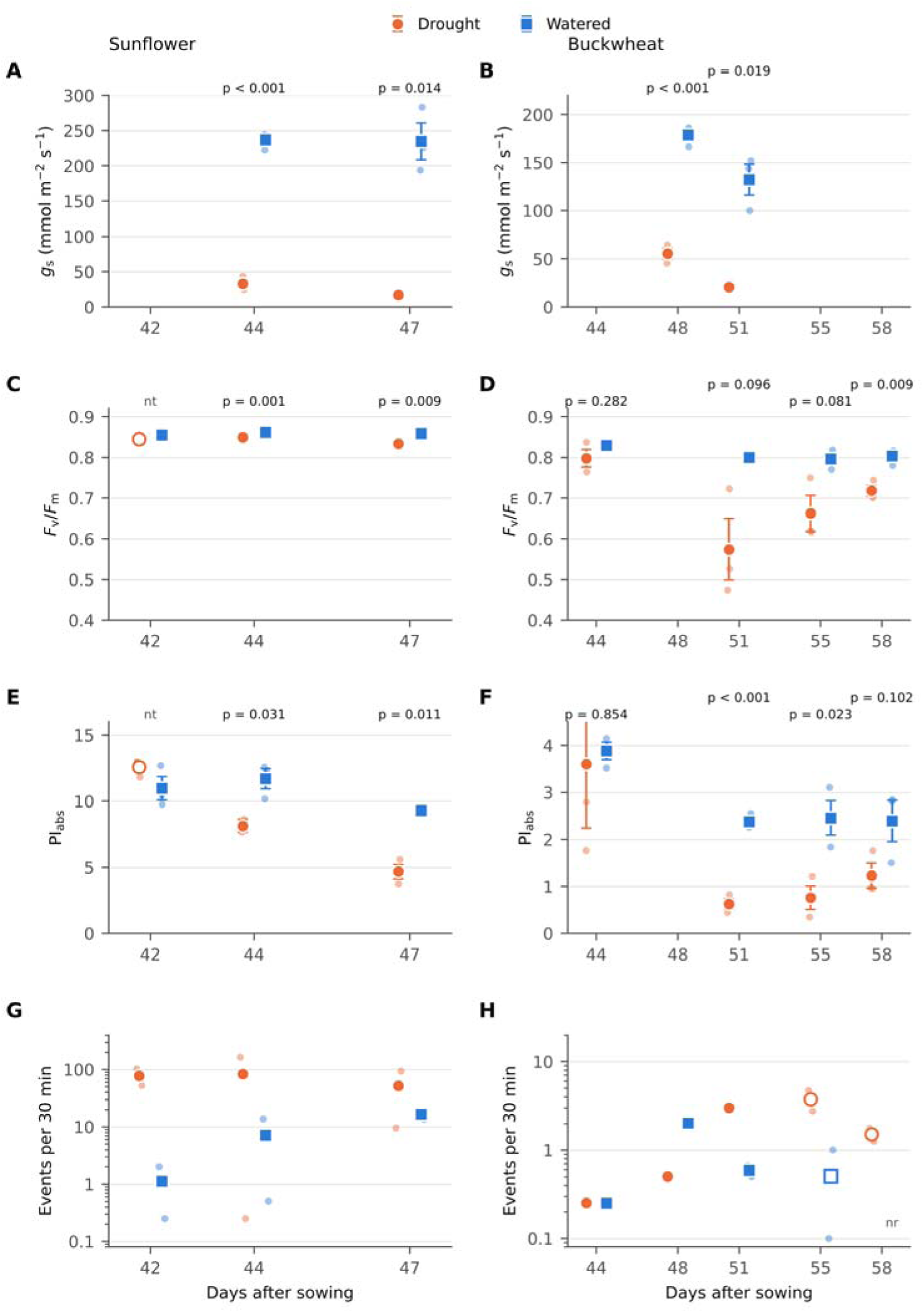
Stomatal conductance (*g*_s_), PSII function and event rates of drought and watered plants. (A-F) Large symbols, mean ± SE of three plants; small dots, single plants, each the mean of two leaves (one gas-exchange reading for one watered buckwheat plant at 51 DAS); p, two-sided Welch’s t-test. Open circle, sunflower drought leaves at 42 DAS with a different background value, not tested (nt); at 44 DAS only leaves with the same background value are included (two drought plants). (G, H) Large symbols, mean of the two end plants; small dots, single end plants; open symbols, raised electrical background; zero rates plotted at 0.1; nr, not recorded. *F*_v_/*F*_m_, maximum quantum yield of PSII; *PI_abs_*, performance index on absorption basis; DAS, days after sowing. **Alt text:** Eight dot plots over the recording days of both species: stomatal conductance far lower in drought plants; maximum PSII yield and performance index of drought sunflower falling only slightly and late, and of drought buckwheat falling strongly from 51 days after sowing; and event rates on a logarithmic scale, higher in drought plants.

Drought lowered PSII performance in both species (Fig. 6C-F; Supplementary Table S7; Supplementary Fig. S2). In sunflower, *F*_v_/*F*_m_ of drought plants stayed at 0.83-0.85. At 42 DAS it was numerically similar to that of watered plants (0.845 against 0.855, not tested), and at 44 and 47 DAS slightly lower (p = 0.001 and 0.009). Their *PI_abs_* was 30% lower than in watered plants at 44 DAS (8.2 against 11.7, n = 2 drought plants, p = 0.031) and half as high at 47 DAS (4.7 against 9.3, p = 0.011). In buckwheat, PI_abs_ of drought plants was similar to that of watered plants at 44 DAS (3.6 against 3.9, p = 0.85) and fell to a quarter at 51 DAS (0.63 against 2.38, p < 0.001), when their *F*_v_/*F*_m_ was 0.57 against 0.80 (p = 0.096). Without two leaves with an *F*_v_/*F*_m_ of 0.20 and 0.32, PI_abs_ was 0.97 (p = 0.001) and *F*_v_/*F*_m_ 0.74 (p = 0.009). *PI_abs_* stayed low at 55 and 58 DAS, and watered buckwheat kept an *F*_v_/*F*_m_ of 0.80-0.83. Over the series, raised event rates coincided with low *g*_s_ in both species (Fig. 6G, H). In sunflower, event rates were already raised at 42 DAS, and a lower *PI_abs_* was detected from 44 DAS. In buckwheat, event rates rose when *PI_abs_* fell.

### Event size and spectrum during drying

The sunflower drought plant at M1 changed its sound as it dried (Fig. 5C, D). The share of its events above −35 dBFS rose from 13% at 42 DAS to 41% at 44 DAS, the day of its highest rate, when its events were 5.7 dB louder (p < 0.001). The power law exponent fell from 1.04 to 0.62, so loud events became more common. At 47 DAS, 5% of its 38 events exceeded −35 dBFS (exponent 1.29). At equal loudness, its share of energy at 40-80 kHz was 24%, 10% and 36% at 42, 44 and 47 DAS (44 against 42 DAS, ratio 0.57, p < 0.001; 47 against 42 DAS, ratio 1.39, p = 0.026), while the watered plant at the same microphone stayed at 36-37%. Within recordings, each 10 dB increase in level reduced this share by a factor of 0.76 (95% CI 0.71-0.82). The drought plant at M2 did not show this relation (factor 1.07, p = 0.49). The summed squared peak amplitude of the plant at M1 rose 9.7 fold from 42 to 44 DAS while its count rose 1.6 fold. Over all drought recordings it was about 710 times higher in sunflower than in buckwheat, against 39 times more events.

The events of both buckwheat drought end plants were pooled (Fig. 5E). They stayed close to the counting level at 51 and 55 DAS (median −49.8 and −49.6 dBFS) and were 3.8 dB louder at 58 DAS (95% CI 1.3-6.4, p = 0.003). Against 55 DAS alone, which had a similar background, the difference was 3.2 dB (p = 0.08).

### Timing of events

Events occurred at random in time (Fig. 5F). In the 15 recordings with at least 30 primary events, all from sunflower drought plants (1,632 events), the coefficient of variation of the intervals between events was 0.85-1.14 (median 0.96), close to the value of 1 expected for random timing. In 14 of the 15 recordings, the Fano factor lay within the range of random simulations at 1, 10 and 60 s. Intervals shorter than 0.25 s were slightly more frequent than expected (38 against 27.8, p = 0.04). The 50 intervals pooled from 17 buckwheat recordings agreed with random timing (Kolmogorov-Smirnov test, p = 0.24).

### Identifying species and drought from the sounds

As a proof of concept, the share of energy at 40-80 kHz alone identified the species of single events from plants not used for training (Fig. 7A, B). Its AUC was 0.91, and 80% of single events were assigned to the correct species. Eight spectral features together did not do better (AUC 0.90). With three events from a left-out plant, the species was correct in 93% of draws, and with five events in 96%, with 85% for the least well classified plant. The 99.6% reached with 10 events is less informative, because the two watered buckwheat plants had only 10 and 13 events. Within each species, the spectrum separated drought and watered plants no better than chance (AUC 0.44 in sunflower and 0.40 in buckwheat), and the event rate of single plant-days separated them with AUCs of 0.76 and 0.70.

**Fig. 7.**
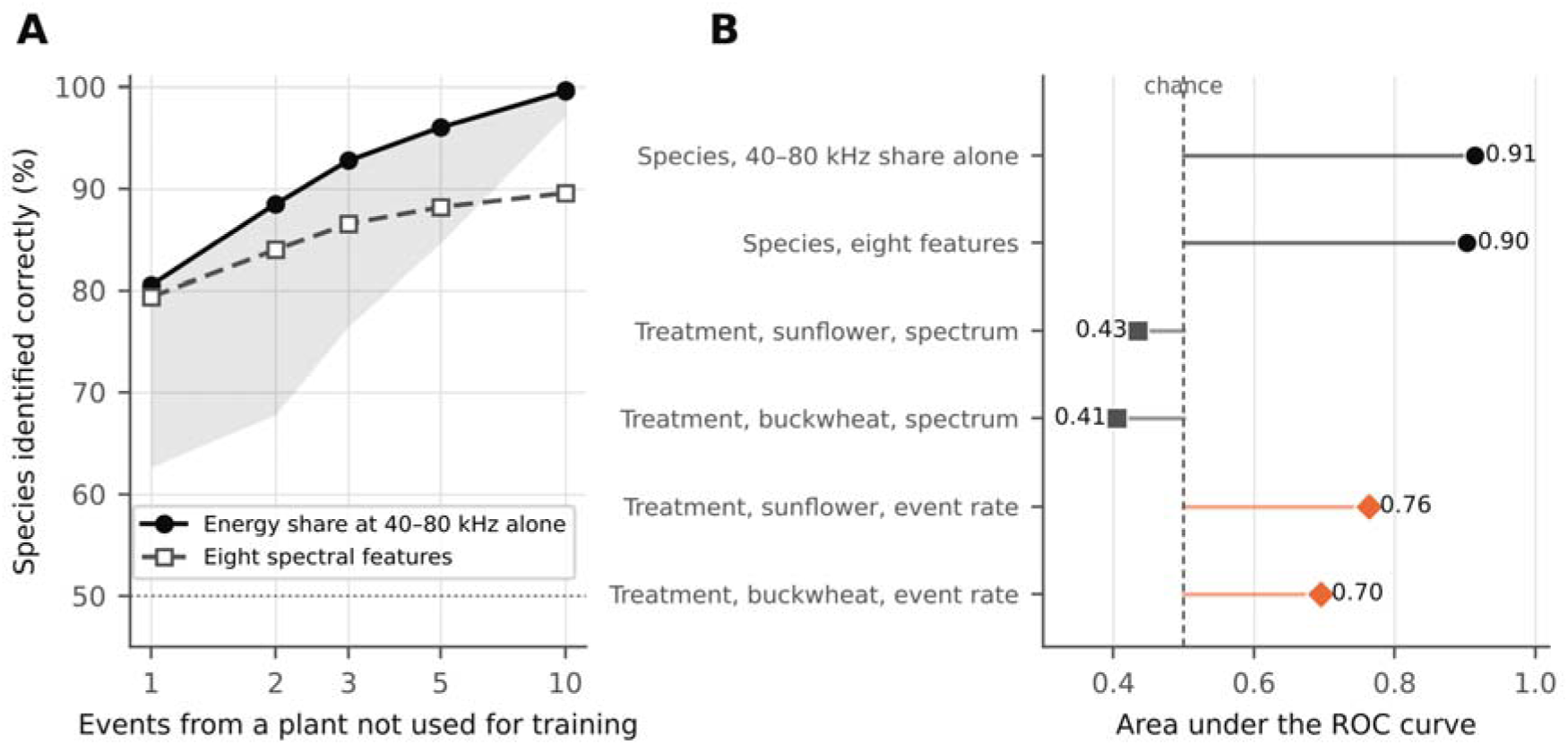
The spectrum identified the species, and the event rate separated drought from watered plants. Logistic classifiers were trained on events between −50.8 and −44 dBFS from seven of the eight end plants and tested on the eighth, for each plant in turn. (A) Mean share of 2,000 random draws of 1-10 events that gave the correct species; shading, range down to the least well classified plant (40-80 kHz share); dotted line, chance. (B) Area under the receiver operating characteristic (ROC) curve for species and treatment from the spectrum of single events, and for treatment from the event rate of single plant-days. **Alt text:** A line plot of species identification accuracy rising with the number of events per plant, and a dot plot of the area under the ROC curve for species and treatment classification.

## Discussion

Our results bear on three points raised by Font Bordera et al. (2026), namely how emission relates to plant water status, whether emissions differ between species and whether classification can support plant monitoring.

### Drought raised emission rates in both species

In sunflower both drought end plants emitted far more than the controls at 42 DAS, and on each later day one of them did (Fig. 2A, B, E; Table 1). In buckwheat both drought plants emitted more from 51 DAS, 14 days after watering stopped (Fig. 2C-E). Absolute rates are hard to compare because thresholds, microphone distance, plant size and counting rules differ (Supplementary Table S8).

We regard 51 DAS as the main evidence for a drought effect in buckwheat. On that day the background was normal and the effect held in every sensitivity analysis (Supplementary Table S2), while the larger ratio at 55 DAS came from recordings with raised background. With two drought and two watered end plants per species, the model estimates are exploratory and their intervals wide (Table 1). The drought effect was clear only on days when both drought plants emitted strongly, and the rate results need confirmation on more plants. Drought plants were also usually recorded in the morning and controls in the early afternoon (Supplementary Table S1), and plant water status changes over the day. The morning slot alone did not produce high rates, because at 48 DAS the afternoon controls of buckwheat emitted more than the morning drought plants, and each sunflower drought plant emitted at or below control level on one morning. Recording the treatments together or alternately would remove this confound. Drought pots were partly rewatered after each recording day, and recovery was not recorded. Sunflower does not refill embolised conduits after rewatering (Stewart et al., 2026), so if events mark embolism, its later events came from conduits not yet embolised.

Emission rate did not simply follow the water content of the substrate (Fig. 5A, B; Supplementary Table S4). The sunflower drought pots dried in parallel, yet the plant at M1 peaked at 44 DAS and nearly stopped by 47 DAS while its pot kept losing water, and the plant at M2 peaked at 47 DAS (Fig. 2A, B). Part of the low at M2 at 44 DAS probably came from the microphone, because the control at M2 was also nearly silent and M2 recorded none of the 275 loud events of the plant at M1. Differences between the plants, in size or xylem vulnerability, are a more likely cause than differences in drying. In both species, emission was highest at intermediate substrate water and lower in the driest pots, as in drying tomato (Khait et al., 2023). A single low rate can therefore come from an unstressed plant or from one past its peak. Control rates in sunflower rose over the series (Fig. 2A, B), probably because the plants grew and their pots dried between waterings, and this narrowed the difference between treatments.

### Emission rose on days of low stomatal conductance

On the days with gas-exchange data, emission was raised only when stomatal conductance of drought plants was low (Fig. 6). In sunflower, at least one drought plant emitted many events on both measurement days, when *g*_s_ of drought plants was 17-33 mmol m^−2^ s^−1^, or 7-14% of that of watered plants. In buckwheat, the event rate of drought plants rose from 0.5 to 3.0 events per 30 min between 48 and 51 DAS, while their *g*_s_ fell from 55 to 21 mmol m^−2^ s^−1^. This association over two days per species does not show that stomatal closure causes the emissions. A simpler explanation is that both changes follow falling water potential. Stomata close at higher water potentials than those that start embolism (Martin-StPaul et al., 2017), and direct observations in leaves showed embolism starting only after stomatal closure (Creek et al., 2020). Closure slows the further rise in xylem tension, but water is still lost through the cuticle and incompletely closed stomata (Duursma et al., 2019), so tension keeps rising as the substrate dries and the most vulnerable conduits cavitate. Emissions are therefore expected mainly after stomatal closure, although two measurement days cannot show exactly when emission began.

In sunflower, emission was already raised on the first recording day, when *F_v_/F_m_* was numerically similar between treatments, and a lower *PI_abs_* was detected from 44 DAS (Fig. 6C, E, G). The fluorescence of that first day could not be tested, only two drought plants could be compared at 44 DAS, and *PI_abs_* is more sensitive to drought than *F*_v_/*F*_m_ (Oukarroum et al., 2007). Our data therefore do not show that emission rose before every change in PSII function. In greenhouse tomato, PSII yield likewise showed no early response while emission did (Dutta et al., 2025). In buckwheat, *PI_abs_* had already fallen to less than half of the control value when emission rose at 51 DAS, with or without the two leaves with very low *F*_v_/*F*_m_ (Fig. 6F, H), which shows that the low emission rate of buckwheat did not come from milder stress.

We propose a conceptual hydraulic model that links these observations (Fig. 8). It is a hypothesis, because we did not measure water potential or embolism. If each event marks one embolising conduit, the event rate is the slope of the vulnerability curve times the rate at which water potential falls. Cumulative emission energy follows the loss of conductivity in drying wood (Mayr and Rosner, 2011), and after stomatal closure water potential falls through residual water loss (Duursma et al., 2019). The model predicts few events before closure, a peak on the steep part of the curve and a decline once the vulnerable conduits are used up. The sunflower plant at M1 followed this course (Fig. 2A), which is consistent with the model without testing it. Sunflower stems lose half their conductivity near −3 MPa (Ahmad et al., 2018), so the events at moderate drying probably came from the most vulnerable conduits. The model could also explain why the two sunflower end plants emitted at rates 10 fold apart at 47 DAS, although both had a *g*_s_ near 18 mmol m^−2^ s^−1^ (Supplementary Table S6). Near its minimum, *g*_s_ no longer tracks water potential, and the two plants could sit at different points on their curves. The plant at M1, past its emission peak, also had the lower PI_abs_ (3.8 against 5.6). Event rate may therefore carry information that gas exchange does not, and calibrating it against leaf water potential, as proposed by Font Bordera et al. (2026), would test the model.

**Fig. 8.**
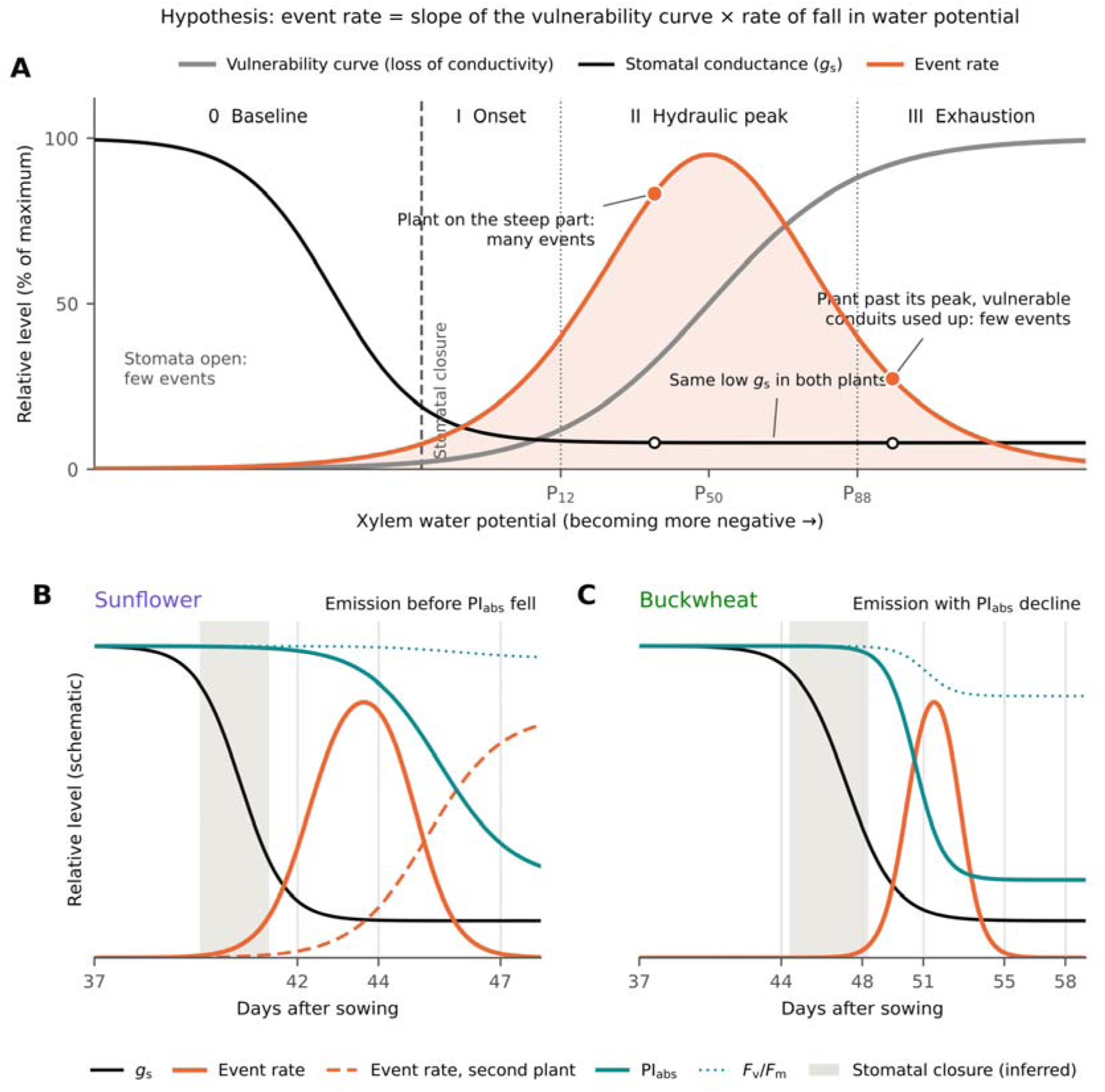
Conceptual hydraulic model, proposed as a hypothesis, linking stomatal closure, xylem embolism, acoustic events and PSII function. (A) Event rate, loss of conductivity and stomatal conductance as xylem water potential falls, if each event marks one embolising conduit. Circles, two plants with the same low stomatal conductance at different points of the curve. P_12_, P_50_ and P_88_, water potentials at 12, 50 and 88% loss of conductivity. (B, C) Hypothesised order of the signals during drying; vertical lines, recording days. Stomatal closure is inferred from wilting at 42 DAS and low *g*_s_ at 44 DAS in sunflower and from *g*_s_ at 48 DAS in buckwheat. Second plant, sunflower end plant at M2. Curves are schematic. DAS, days after sowing; *g*_s_, stomatal conductance; *PI_abs_*, performance index on absorption basis; *F*_v_/*F*_m_, maximum quantum yield of PSII. **Alt text:** Schematic diagrams: a sigmoid vulnerability curve with a bell-shaped event-rate curve that peaks on its steep part, with stomatal closure marked before embolism starts; and two timelines in which stomatal conductance falls first and the event rate then rises, before the PSII performance index falls in sunflower and together with its fall in buckwheat.

### The two species differ in the spectral content of their events

At equal loudness, sunflower events carried about 10 times more of their energy at 40-80 kHz than buckwheat events (Fig. 3C; Table 2), and about half of them reached the 60-80 kHz band, against none in buckwheat (Fig. 3D; Supplementary Table S3). The difference held in every check and did not differ between drought and watered plants (Fig. 3C, D). Most events of both species peaked near 29-30 kHz (Fig. 3B; Table 2). A peak shared by two species probably reflects the response of the microphone and chamber, because the waveform changes on its way from the source to the sensor (Vergeynst et al., 2015), and microphones near tree saplings also recorded about 30 kHz on average (Klaminder et al., 2025). Khait et al. (2023) separated tomato and tobacco by mean peak frequency. In our data peak frequency was a weak marker, and the energy above 40 kHz and band specific detection, which a shared resonance affects less, were much stronger. Font Bordera et al. (2026) described plant emissions as specific to both species and stress, and our data support species specificity for sunflower and buckwheat. At equal loudness, the spectrum did not differ between drought and watered plants over all days, although it shifted during drying in one plant (Fig. 5D).

The difference is a property of the plants and the recording geometry together, and our data cannot say which plant property produces it. The microphones, chamber, thresholds and microphone-to-stem distance were the same for both species, and the matched loudness removes differences in event level. The species were, however, recorded in separate series, buckwheat in the two weeks after sunflower, so a change in the recording chain between the series cannot be excluded, although the difference held on every recording day of both series. Three plant-related sources remain. The first is the source itself. Font Bordera et al. (2026) attributed species signatures to vessel diameter, tissue anatomy or wood density, and in the vessel model of Dutta et al. (2022) resonance frequency falls and settling time rises with vessel radius. A source origin would therefore require narrower vessels in sunflower, whose events were shorter and richer in high frequencies than those of buckwheat. The second is the transmission of the event through stem and leaf tissue, which depends on stem diameter and on tissue properties, because lignified xylem walls carry high frequencies and softer tissues damp them (Font Bordera et al., 2026). The third is radiation into the air, which depends on plant size and on where the emitting tissue sits relative to the microphone. The taller sunflower plants would have placed more sources further from the microphone, which would lower their high-frequency share. Vessel diameters, a contact sensor on the stem recording with the microphone, and recordings at several distances would separate these sources.

### Classification as a proof of concept

The share at 40-80 kHz identified the species of plants not used for training (Fig. 7A). Single events were assigned correctly in 80% of cases, similar to the 70-80% reported for single tomato and tobacco sounds (Khait et al., 2023), and five events from a plant gave at least 85% accuracy. Font Bordera et al. (2026) presented classification of plant ultrasound as a route to precision agriculture. Our result comes from four plants per species of one cultivar in one chamber, and is a proof of concept. A diagnostic method would need validation on many more plants, growth stages and recording systems. It does show that one interpretable spectral feature can carry most of the species’ information. The spectrum did not separate drought and watered plants, while the event rate did (Fig. 7B). A sensor could use the spectrum to identify the crop and the course of the event rate to detect drought, because a plant past its emission peak can emit as little as an unstressed one. A monitoring system would also need a sensitivity check for each microphone, because a failing microphone looks like a quiet plant; the events recorded by two microphones in common provide one.

### Shared events suggest that each microphone recorded mainly the nearest plant

At 343 m s^−1^, the 2.24 ms difference in arrival time between the two groups of shared events (Fig. 4B) corresponds to a path difference of about 0.38 m, which fits a source near one end of the row. With the near microphone 5 cm from the stem, spherical spreading to the far microphone, about 0.43 m away, predicts a loss of 19 dB, close to the observed 21 dB (Fig. 4A). Events from the middle plant would arrive about 14 dB weaker, and only 27% of sunflower and 1% of buckwheat events would still be detected. Shared events were few, however (35 of 621 events at 42 DAS) and came mostly from one day, so events from the middle plant cannot be excluded, and we treat M1 and M2 as positions dominated by the nearest plant. The far copies of events from the plant at M1 lost more than half of their high frequency share (Fig. 4C), although the extra distance was the same in both directions. This points to the microphones or their orientation, which studies comparing spectra should keep fixed.

### The size and spectrum of events changed as plants dried

In the sunflower plant at M1, events at the emission peak were louder and carried less energy above 40 kHz than two days earlier, and after the peak the few remaining events were quiet and rich in high frequencies (Fig. 5C, D). Within recordings, louder events also had less high-frequency energy. More distant sources would give quieter events with less high-frequency energy, the opposite relation, and the watered plant at the same microphone had stable spectra, which places the relation in the plant. If loud, low-pitched events come from large conduits (Mayr and Rosner, 2011; Ponomarenko et al., 2014; Khait et al., 2023), this plant would have lost mainly large conduits at its emission peak and mainly small ones later. The amplitude and frequency of emissions do not always reflect embolism size, however (Vergeynst et al., 2015; Lamacque et al., 2022; Font Bordera et al., 2026), and this interpretation rests on one plant. The summed squared peak amplitude amplified the changes in count, as expected if emission energy follows the loss of conductivity (Mayr and Rosner, 2011).

### Events occurred at random in time

Fast cascades of embolism spreading between conduits would cluster events at short intervals, but events occurred at random in time (Fig. 5F). We found only about 10 extra pairs shorter than 0.25 s among 1,617 intervals, so most detected events were independent. With random timing, a count of 100 events gives the rate within about ±20%, which would take about 25 h at the buckwheat rate of about two events per 30 min.

## Conclusions

Airborne ultrasonic emissions of sunflower and buckwheat carried species identity in their spectrum and drought in their event rate. At equal loudness, sunflower events carried about 10 times more energy at 40-80 kHz than buckwheat events in drought and watered plants alike, although events of both species peaked near 30 kHz. As a proof of concept on four plants per species and one cultivar, this share identified the species from a few events. Every drought plant emitted more events than the watered plants of its species. In single plants the rate rose and then fell, so a low rate can come from an unstressed plant or from one past its emission peak. The rise coincided with low stomatal conductance, as expected if emissions arise as xylem tension keeps rising after stomata close, and in sunflower emission was already raised when maximum PSII quantum yield was still similar between treatments. To our knowledge, these are the first airborne recordings of either species. Recordings of more plants, with both treatments at the same time of day and with leaf water potential, stem anatomy and contact sensors on the same plants, will be needed to link these signals to plant hydraulics.

## Author contributions

BS: conceptualization, investigation, data curation, formal analysis, methodology, writing - original draft, writing - review and editing; MIH: investigation, data curation, formal analysis, writing - review and editing; MS: investigation; AR: conceptualization, supervision, writing - review and editing, resources.

## Use of generative AI

During the preparation of this work, the author used the AI-assisted tool Claude (Anthropic) to improve the readability and language of the manuscript and to assist with drafting figures. The author reviewed and edited all content and takes full responsibility for the publication.

## Data availability

The recordings, event tables and analysis scripts (R and Python) are available from the corresponding author upon request.

## Abbreviations

AE: acoustic emission
AUC: area under the receiver operating characteristic curve
CI: confidence interval
DAS: days after sowing
dBFS: decibels relative to full scale
*F*_v_/*F*_m_: maximum quantum yield of PSII
*g*_s_: stomatal conductance to water vapour
*PI_abs_*: performance index on absorption basis

