## Supplementary Data for "Ultrasonic emissions of sunflower and buckwheat carry species identity in their spectrum and drought in their rate"

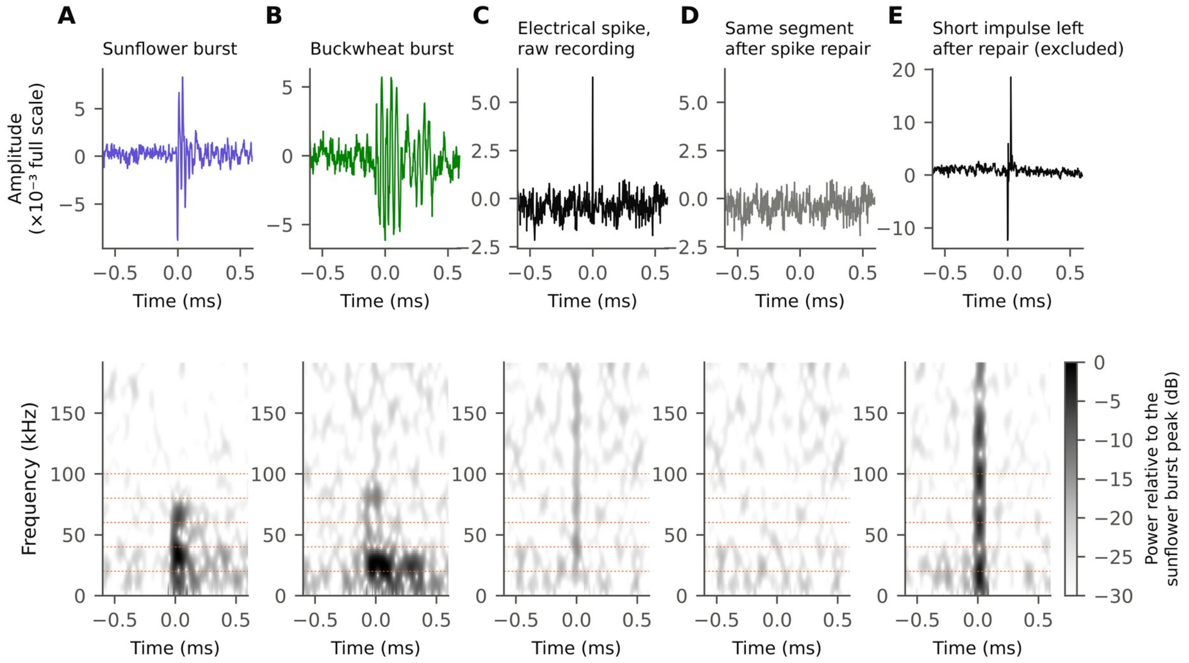


**Fig. S1.** Acoustic bursts compared with electrical spikes. Top, waveforms; bottom, spectrograms (0.125-ms Hann windows). (A) Sunflower drought plant (47 DAS, microphone M2, −46 dBFS). (B) Buckwheat drought plant (55 DAS, microphone M1, −46 dBFS). (C, D) One-sample spike from the mains USB power supply before and after repair (watered buckwheat, 44 DAS, M2). (E) Impulse excluded by the waveform check (drought buckwheat, 44 DAS, M2). Dashed lines, band edges. DAS, days after sowing.

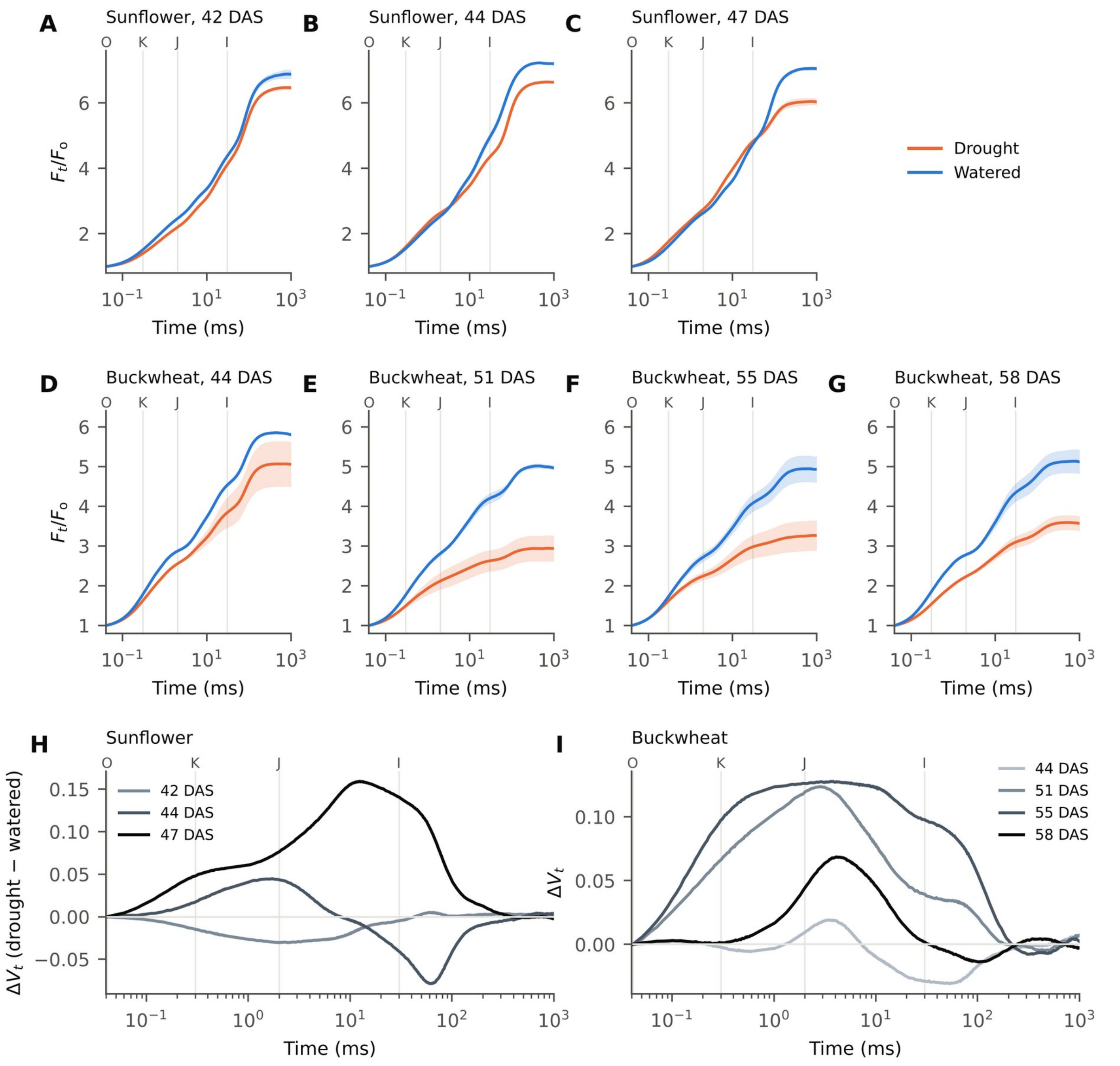


**Fig. S2.** Chlorophyll *a* fluorescence transients of drought and watered plants. (A-G) Fluorescence relative to initial fluorescence ($F_t/F_o$); lines, mean of three plants (two drought plants at 44 DAS), each the mean of two leaves; shading, SE. (H, I) Difference in relative variable fluorescence, $V_t = (F_t - F_o)/(F_m - F_o)$, between drought and watered plants. O, K, J and I, 0.04, 0.3, 2 and 30 ms. DAS, days after sowing; $F_m$, maximum fluorescence.

**Table S1.** Recording schedule and electrical background of each group of recordings

| Species | DAS | Treatment | Mic | n | Time window | Background ÷ empty chamber | Repaired spike samples per s |
|---|---|---|---|---|---|---|---|
| Sunflower | 42 | Drought | M1 | 4 | 10:35-12:35 | 1.03-1.04 | 57-65 |
| Sunflower | 42 | Drought | M2 | 4 | 10:35-12:35 | 1.06-1.07 | 62-69 |
| Sunflower | 42 | Watered | M1 | 4 | 13:40-15:40 | 1.02-1.02 | 53-60 |
| Sunflower | 42 | Watered | M2 | 4 | 13:40-15:40 | 1.02-1.06 | 54-107 |
| Sunflower | 44 | Drought | M1 | 4 | 10:15-12:15 | 1.01-1.02 | 38-39 |
| Sunflower | 44 | Drought | M2 | 4 | 10:15-12:15 | 1.04-1.09 | 42-81 |
| Sunflower | 44 | Watered | M1 | 4 | 12:40-14:40 | 1.02-1.03 | 50-58 |
| Sunflower | 44 | Watered | M2 | 4 | 12:40-14:40 | 1.02-1.04 | 58-95 |
| Sunflower | 47 | Drought | M1 | 4 | 10:10-12:10 | 1.05-1.06 | 100-105 |
| Sunflower | 47 | Drought | M2 | 4 | 10:10-12:10 | 1.01-1.02 | 51-54 |
| Sunflower | 47 | Watered | M1 | 4 | 12:40-14:40 | 1.00-1.02 | 32-33 |
| Sunflower | 47 | Watered | M2 | 4 | 12:40-14:40 | 1.01-1.03 | 44-45 |
| Buckwheat | 44 | Drought | M1 | 4 | 11:15-13:15 | 1.01-1.01 | 37-38 |
| Buckwheat | 44 | Drought | M2 | 4 | 14:20-16:20 | 1.10-1.14 | 90-130 |
| Buckwheat | 44 | Watered | M1 | 4 | 14:20-16:20 | 1.00-1.01 | 32-37 |
| Buckwheat | 44 | Watered | M2 | 4 | 11:15-13:15 | 1.15-1.17 | 168-188 |
| Buckwheat | 48 | Drought | M1 | 4 | 11:00-13:00 | 1.04-1.06 | 79-87 |
| Buckwheat | 48 | Drought | M2 | 4 | 11:00-13:00 | 1.02-1.04 | 38-39 |
| Buckwheat | 48 | Watered | M1 | 4 | 14:00-16:00 | 1.04-1.05 | 57-72 |
| Buckwheat | 48 | Watered | M2 | 4 | 14:00-16:00 | 1.04-1.04 | 42-45 |
| Buckwheat | 51 | Drought | M1 | 4 | 11:10-13:10 | 1.02-1.02 | 40-42 |
| Buckwheat | 51 | Drought | M2 | 4 | 11:10-13:10 | 1.03-1.04 | 80-86 |
| Buckwheat | 51 | Watered | M1 | 3 | 13:45-15:15 | 1.03-1.03 | 56-59 |
| Buckwheat | 51 | Watered | M2 | 4 | 13:45-15:45 | 1.01-1.04 | 46-57 |
| Buckwheat | 55 | Drought | M1 | 4 | 11:05-13:05 | 1.28-1.33 | 160-170 |
| Buckwheat | 55 | Drought | M2 | 4 | 11:05-13:05 | 1.24-1.25 | 187-197 |
| Buckwheat | 55 | Watered | M1 | 4 | 13:35-15:35 | 1.12-1.15 | 34-166 |
| Buckwheat | 55 | Watered | M2 | 4 | 13:35-15:35 | 1.23-1.25 | 110-128 |
| Buckwheat | 58 | Drought | M1 | 4 | 13:25-15:25 | 1.27-1.28 | 138-185 |
| Buckwheat | 58 | Drought | M2 | 4 | 13:25-15:25 | 1.22-1.23 | 89-92 |

Mic, microphone next to the end plant; n, number of 30-min recordings; time window, start of the first to end of the last recording; background ÷ empty chamber, background level relative to the empty-chamber recordings used to set the thresholds (above 1.1, raised electrical background); repaired spike samples per s, range across recordings. DAS, days after sowing.

**Table S2.** Sensitivity of the buckwheat drought effect on event rates to the recordings with raised electrical background

| Data set | Recordings | Treatment × date (likelihood-ratio $\chi^2$, df, p) | Rate ratio at 51 DAS (95% CI), p | Rate ratio at 55 DAS (95% CI), p | Rate ratio, 51 and 55 DAS pooled (95% CI) |
|---|---|---|---|---|---|
| All recordings | 63 | 23.0, 3, < 0.001 | 5.3 (1.8-15.1), 0.002 | 7.5 (2.6-21.3), < 0.001 | 6.4 (3.0-13.4) |
| Without 55 DAS | 47 | 16.9, 2, < 0.001 | 5.3 (1.8-15.1), 0.002 | - | - |
| Normal background only | 39 | 16.5, 2, < 0.001 | 5.3 (1.8-15.1), 0.002 | - | - |
| 51 DAS alone | 15 | - | 5.3 (1.8-15.1), 0.002 | - | - |
| All recordings, background ratio as covariate | 63 | - | 5.3 (1.8-15.2), 0.002 | 4.5 (1.3-14.8), 0.015 | - |

Rate ratios (drought/watered) from Poisson generalised linear mixed models with treatment × date, random intercepts for plant and plant-day and an offset for recording length (51 DAS alone, random intercept for plant); p values are two-sided. Recordings at 58 DAS had no watered controls and are not included. Normal background only excludes the recordings at 55 DAS and the M2 recordings at 44 DAS. Background ratio, background level of each recording relative to the empty chamber. DAS, days after sowing; -, not estimated.

**Table S3.** Band-specific detection of ultrasonic events from sunflower and buckwheat by treatment and plant

| Species | Treatment | Plant | n | Median level (dBFS) | 20-40 kHz (%) | 40-60 kHz (%) | 60-80 kHz (%) | 80-100 kHz (%) | Two or more bands (%) | Strongest band above 40 kHz (%) |
|---|---|---|---|---|---|---|---|---|---|---|
| **Events between −50.8 and −44 dBFS, recordings with normal background** | | | | | | | | | | |
| Sunflower | Drought | M1 | 345 | −47.5 | 93.0 | 82.9 | 32.2 | 0.3 | 80.9 | 11.6 |
| Sunflower | Drought | M2 | 432 | −48.3 | 85.0 | 81.0 | 60.6 | 0.9 | 82.4 | 21.1 |
| Sunflower | Watered | M1 | 96 | −48.0 | 81.2 | 81.2 | 53.1 | 3.1 | 81.2 | 31.2 |
| Sunflower | Watered | M2 | 42 | −48.9 | 85.7 | 66.7 | 47.6 | 19.0 | 69.0 | 23.8 |
| Buckwheat | Drought | M1 | 15 | −49.7 | 100.0 | 6.7 | 0.0 | 0.0 | 6.7 | 0.0 |
| Buckwheat | Drought | M2 | 14 | −50.3 | 100.0 | 0.0 | 0.0 | 0.0 | 0.0 | 0.0 |
| Buckwheat | Watered | M1 | 10 | −49.2 | 100.0 | 10.0 | 0.0 | 0.0 | 10.0 | 0.0 |
| Buckwheat | Watered | M2 | 8 | −47.2 | 100.0 | 0.0 | 0.0 | 0.0 | 0.0 | 0.0 |
| **All primary events (above −50.8 dBFS), all recordings** | | | | | | | | | | |
| Sunflower | Drought | Both | 1698 | −43.1 | 94.4 | 91.0 | 56.4 | 3.4 | 91.1 | 13.7 |
| Sunflower | Watered | Both | 198 | −46.9 | 87.4 | 79.3 | 56.6 | 14.1 | 79.8 | 28.8 |
| Buckwheat | Drought | Both | 72 | −49.5 | 98.6 | 12.5 | 2.8 | 1.4 | 12.5 | 1.4 |
| Buckwheat | Watered | Both | 26 | −47.9 | 100.0 | 15.4 | 7.7 | 0.0 | 15.4 | 7.7 |

Shares of events above the detection threshold in each band, detected in two or more bands and strongest in a band above 40 kHz. M1 and M2, end plants next to microphone M1 or M2; both, both end plants; n, number of events. Normal background excludes the buckwheat recordings at 55 and 58 DAS and the M2 recordings at 44 DAS. DAS, days after sowing.

**Table S4.** Pot weights and substrate water of the drought and watered pots of sunflower and buckwheat

| DAS | Plant | Drought pot before rewatering (g) | Drought pot after rewatering (g) | Substrate water before rewatering, upper limit (% of field capacity) | Water lost by the drought pot per day until the next weighing (g) | Watered pot (g) |
|---|---|---|---|---|---|---|
| **Sunflower** | | | | | | |
| 42 | M1 | 222 | 237 | 60 | 36.0 | 248 |
| 42 | M2 | 195 | 210 | 50 | 20.0 | 277 |
| 42 | Middle | 223 | 230 | 61 | 33.0 | 317 |
| 43 | M1 | 201 | 213 | 52 | 19.8 | 322 |
| 43 | M2 | 190 | 201 | 48 | 17.4 | 341 |
| 43 | Middle | 197 | 209 | 51 | 23.6 | 332 |
| 44 | M1 | 193 | 209 | 49 | 21.3 | 271 |
| 44 | M2 | 184 | 203 | 46 | 21.0 | 278 |
| 44 | Middle | 185 | 212 | 46 | 23.7 | 283 |
| 47 | M1 | 165 | - | 38 | - | 347 |
| 47 | M2 | 160 | - | 36 | - | 401 |
| 47 | Middle | 161 | - | 37 | - | 385 |
| **Buckwheat** | | | | | | |
| 44 | M1 | 261 | 286 | 75 | 17.5 | 364 |
| 44 | M2 | 268 | 283 | 78 | 19.5 | 332 |
| 44 | Middle | 251 | 285 | 72 | 14.8 | 374 |
| 48 | M1 | 216 | 240 | 58 | 10.3 | 408 |
| 48 | M2 | 205 | 237 | 54 | 12.3 | 405 |
| 48 | Middle | 226 | 238 | 62 | 13.3 | 413 |
| 51 | M1 | 209 | - | 55 | 6.0 | 456 |
| 51 | M2 | 200 | - | 52 | 5.2 | 442 |
| 51 | Middle | 198 | - | 51 | 6.0 | 431 |
| 55 | M1 | 185 | - | 46 | 10.0 | 440 |
| 55 | M2 | 179 | - | 44 | 5.0 | 438 |
| 55 | Middle | 174 | - | 42 | 4.3 | 460 |
| 58 | M1 | 155 | - | 34 | - | 461 |
| 58 | M2 | 164 | - | 38 | - | 454 |
| 58 | Middle | 161 | - | 37 | - | 472 |

Pot weights include the pot (18.77 g), substrate and plant and were taken after recording; 43 DAS was a weighing day without recordings. Substrate water = (pot weight − 66.5 g) ÷ 258 g × 100, an upper limit because plant mass was not subtracted. Drought pots were rewatered to about 40-50% of field capacity (sunflower, 42-44 DAS; buckwheat, 44 and 48 DAS); water lost from 44 to 47 DAS includes about 10 mL added to each sunflower drought pot at 45 and 46 DAS. Watered pots were watered every two days. M1 and M2, end plants next to microphone M1 or M2; DAS, days after sowing; -, not rewatered.

**Table S5.** Stomatal conductance, transpiration, intercellular $CO_2$ and PSII quantum yield of sunflower and buckwheat drought and watered plants

| **Species** | **DAS** | **Treatment** | **$g_s$ (mmol m$^{-2}$ s$^{-1}$)** | **$E$ (mmol m$^{-2}$ s$^{-1}$)** | **$c_i$ (µmol mol$^{-1}$)** | **ΦPSII** |
|---|---|---|---|---|---|---|
| Sunflower | 44 | Drought | 33 ± 6 | 0.31 ± 0.05 | 399 ± 4 | 0.162 ± 0.010 |
| | | Watered | 237 ± 7 | 1.85 ± 0.11 | 348 ± 8 | 0.219 ± 0.060 |
| | | *p* | < 0.001 | 0.001 | 0.012 | 0.448 |
| | 47 | Drought | 17 ± 1 | 0.17 ± 0.02 | 477 ± 29 | 0.138 ± 0.009 |
| | | Watered | 235 ± 26 | 1.77 ± 0.14 | 342 ± 3 | 0.211 ± 0.053 |
| | | *p* | 0.014 | 0.007 | 0.043 | 0.302 |
| Buckwheat | 48 | Drought | 55 ± 6 | 0.65 ± 0.06 | 369 ± 2 | 0.166 ± 0.022 |
| | | Watered | 179 ± 6 | 1.63 ± 0.05 | 387 ± 2 | 0.072 ± 0.007 |
| | | *p* | < 0.001 | < 0.001 | 0.005 | 0.039 |
| | 51 | Drought | 21 ± 2 | 0.23 ± 0.02 | 385 ± 10 | 0.145 ± 0.021 |
| | | Watered | 132 ± 16 | 1.28 ± 0.12 | 375 ± 3 | 0.086 ± 0.022 |
| | | *p* | 0.019 | 0.011 | 0.432 | 0.126 |

Mean ± SE of three plants, each the mean of two leaves (one leaf for one watered buckwheat plant at 51 DAS). $g_s$, stomatal conductance to water vapour; $E$, transpiration; $c_i$, intercellular $CO_2$ concentration; ΦPSII, effective quantum yield of PSII; $p$, two-sided Welch's t-test; DAS, days after sowing.

**Table S6.** Stomatal conductance, transpiration and event rate of single sunflower and buckwheat plants at each position in the row

| Species | DAS | Treatment | Position | $g_s$ (mmol m$^{-2}$ s$^{-1}$) | $E$ (mmol m$^{-2}$ s$^{-1}$) | Events per 30 min |
|---|---|---|---|---|---|---|
| Sunflower | 44 | Drought | M1 | 30 | 0.28 | 165.9 |
| | | | M2 | 25 | 0.23 | 0.25 |
| | | | Middle | 44 | 0.41 | - |
| | | Watered | M1 | 223 | 1.64 | 13.8 |
| | | | M2 | 244 | 1.89 | 0.50 |
| | | | Middle | 244 | 2.02 | - |
| | 47 | Drought | M1 | 19 | 0.20 | 9.5 |
| | | | M2 | 17 | 0.16 | 93.9 |
| | | | Middle | 16 | 0.15 | - |
| | | Watered | M1 | 226 | 1.67 | 19.5 |
| | | | M2 | 284 | 2.05 | 13.5 |
| | | | Middle | 194 | 1.59 | - |
| Buckwheat | 48 | Drought | M1 | 56 | 0.67 | 0.50 |
| | | | M2 | 65 | 0.75 | 0.50 |
| | | | Middle | 46 | 0.55 | - |
| | | Watered | M1 | 187 | 1.62 | 2.0 |
| | | | M2 | 182 | 1.72 | 2.0 |
| | | | Middle | 167 | 1.55 | - |
| | 51 | Drought | M1 | 18 | 0.21 | 3.0 |
| | | | M2 | 25 | 0.27 | 3.0 |
| | | | Middle | 19 | 0.20 | - |
| | | Watered | M1 | 152 | 1.45 | 0.67 |
| | | | M2 | 100 | 1.04 | 0.50 |
| | | | Middle | 144 | 1.36 | - |

$g_s$ and $E$, means of two leaves per plant (one leaf for the watered buckwheat plant at M2 at 51 DAS); events per 30 min, means of four 30-min recordings (three for the watered buckwheat plant at M1 at 51 DAS). M1 and M2, end plants next to microphone M1 or M2; middle, plant between them, whose events were not counted separately. $g_s$, stomatal conductance to water vapour; $E$, transpiration; DAS, days after sowing.

**Table S7.** Chlorophyll fluorescence parameters of sunflower and buckwheat drought and watered plants

| Species | DAS | Treatment | $F_v/F_m$ | $PI_{abs}$ | $V_J$ | $V_I$ |
|---|---|---|---|---|---|---|
| Sunflower | 42[a] | Drought | 0.845 ± 0.002 | 12.57 ± 0.36 | 0.219 ± 0.000 | 0.568 ± 0.007 |
| | | Watered | 0.855 ± 0.003 | 10.97 ± 0.88 | 0.249 ± 0.005 | 0.572 ± 0.012 |
| | | *p* | - | - | - | - |
| | 44[b] | Drought | 0.849 ± 0.001 | 8.16 ± 0.50 | 0.290 ± 0.007 | 0.594 ± 0.007 |
| | | Watered | 0.862 ± 0.000 | 11.69 ± 0.76 | 0.247 ± 0.009 | 0.634 ± 0.020 |
| | | *p* | 0.001 | 0.031 | 0.031 | 0.176 |
| | 47 | Drought | 0.834 ± 0.003 | 4.67 ± 0.53 | 0.346 ± 0.017 | 0.757 ± 0.003 |
| | | Watered | 0.858 ± 0.001 | 9.29 ± 0.10 | 0.270 ± 0.002 | 0.616 ± 0.009 |
| | | *p* | 0.009 | 0.011 | 0.046 | 0.002 |
| Buckwheat | 44 | Drought | 0.798 ± 0.021 | 3.60 ± 1.36 | 0.396 ± 0.050 | 0.698 ± 0.007 |
| | | Watered | 0.829 ± 0.001 | 3.88 ± 0.19 | 0.385 ± 0.008 | 0.726 ± 0.016 |
| | | *p* | 0.282 | 0.854 | 0.848 | 0.210 |
| | 51[c] | Drought | 0.574 ± 0.076 | 0.63 ± 0.11 | 0.649 ± 0.030 | 0.880 ± 0.030 |
| | | Watered | 0.800 ± 0.002 | 2.38 ± 0.09 | 0.451 ± 0.008 | 0.797 ± 0.039 |
| | | *p* | 0.096 | < 0.001 | 0.016 | 0.173 |
| | 55 | Drought | 0.662 ± 0.044 | 0.76 ± 0.25 | 0.602 ± 0.059 | 0.888 ± 0.023 |
| | | Watered | 0.796 ± 0.014 | 2.46 ± 0.36 | 0.436 ± 0.009 | 0.778 ± 0.012 |
| | | *p* | 0.081 | 0.023 | 0.102 | 0.023 |
| | 58 | Drought | 0.718 ± 0.014 | 1.23 ± 0.27 | 0.480 ± 0.020 | 0.807 ± 0.003 |
| | | Watered | 0.803 ± 0.011 | 2.39 ± 0.44 | 0.435 ± 0.024 | 0.810 ± 0.010 |
| | | *p* | 0.009 | 0.102 | 0.221 | 0.747 |

Mean ± SE of three plants, each the mean of two leaves. $F_v/F_m$, maximum quantum yield of PSII; $PI_{abs}$, performance index on absorption basis; $V_J$ and $V_I$, relative variable fluorescence at 2 and 30 ms; *p*, two-sided Welch's t-test; DAS, days after sowing; -, not tested. [a] Drought leaves with a different background value, not tested; one watered leaf with that value is not included. [b] Leaves with the same background value only (two drought plants). [c] Includes two drought leaves with $F_v/F_m$ of 0.20 and 0.32; without them, $F_v/F_m$ was 0.74 ($p$ = 0.009) and $PI_{abs}$ 0.97 ($p$ = 0.001).

**Table S8.** Airborne recordings of plant ultrasonic emissions with microphones in this and earlier studies, by plant material, set-up, detection, event rate and spectral measure

| **Study** | **Plants, stress and setting** | **Microphone and distance** | **Detection** | **Event rate** | **Spectral measure** |
|---|---|---|---|---|---|
| Khait et al. (2023) | Tomato and tobacco; drought and cutting; acoustic box (50 × 100 × 150 cm) with three plants | CM16 condenser microphones (Avisoft), two directional microphones per plant, 10 cm | 500 kHz; 20 kHz high-pass; recording triggered above 2% of the microphone dynamic range | Drought: 35.4 $h^{-1}$ (tomato) and 11.0 $h^{-1}$ (tobacco); controls < 1 $h^{-1}$; pots with soil, none in > 500 h | Mean peak frequency 49.6 and 54.8 kHz in drought plants; classifier separated species and treatments |
| Bonisoli et al. (2024) | Pinto bean, unstressed; indoor room, four plants per session; soil-only and empty-room controls | Ultramic 384K BLE (Dodotronic), 1-2 cm from the closest stem | 384 kHz; threshold per band (20-40, 40-60, 60-80, 80-100 kHz) from the background level | Protocol study; rates not compared between treatments | Four 20-kHz bands |
| Bonisoli et al. (2025) | Pinto bean and tomato, 4 weeks old; water stress; indoor and outdoor | Ultramic 384K BLE (Dodotronic), protocol of Bonisoli et al. (2024) | Threshold per band (20-40 to 80-100 kHz) | 2.83 ± 0.49 per 30 min in water-stressed plants, more than in controls | Four 20-kHz bands |
| Dutta et al. (2022) | Shoots of 10 species, cut and dried on the bench; laboratory | M500-USB (Pettersson; 10-150 kHz), about 2 mm from the cut face (axial) and beside the stem (radial) | Single pulses | - | Settling time and resonance frequency, related to vessel radius |
| Dutta et al. (2025) | High-wire tomato in a greenhouse; irrigation stopped | M500-USB (Pettersson), close to the stem in a casing with acoustic foam | Triggered above a 16 kHz cut-off | Drought plants 1,196 and 185 events, controls 77 and 15, over the drought period | Resonance frequency |
| Klaminder et al. (2025) | Cut birch and Scots pine saplings, embolism induced with a surfactant; outdoors | CM16/CMPA-P48 (Avisoft), < 3 cm from the lower trunk, with contact microphones on the trunk | 192 kHz; pulses at 20-65 kHz above a prominence threshold | 58 ± 28% more pulses in treated than control birch; the distant microphone recorded about 82% of the contact counts | Average frequency about 30 kHz in both species and treatments |
| This study | Sunflower and buckwheat; water withheld from 37 DAS; wooden chamber lined with acoustic foam, three plants per row | Ultramic 384K BLE (Dodotronic), one at each end of the row, 5 cm from the nearest stem | 384 kHz; band thresholds 1 dB above empty-chamber maxima; events above −50.8 dBFS | End plants, per 30 min: sunflower drought 0.25-166, watered 0.25-19.5; buckwheat drought 0.25-4.8, watered 0-2.0 | Share of energy at 40-80 kHz (24% sunflower, 2% buckwheat); band detection; peak frequency near 30 kHz in both |

Rates are given as reported. Thresholds, distances and counting rules differ between studies, so absolute rates are not directly comparable. DAS, days after sowing; -, not reported.
